# Electrostatic Self-Assembly of Protein Nanoparticles from Intrinsically Disordered Polypeptide and Globular Protein Pairs

**DOI:** 10.64898/2026.01.07.698192

**Authors:** So Yeon Ahn, Megan S. K. Jen, Dayoung Gloria Lee, Jake D. Johnston, Jane Liao, Eunsol Kim, Ying Tong Yue, Shana Elbaum-Garfinkle, Oleg Gang, Allie C. Obermeyer

## Abstract

Intracellular delivery of folded, functional proteins remains a critical barrier to realizing the full potential of protein therapeutics. We present a fully aqueous, genetically encoded platform for assembling micellar protein nanoparticles via electrostatic coacervation between anionic globular proteins and multi-domain intrinsically disordered proteins (IDPs) composed of a neutral elastin-like polypeptide (ELP) domain and a cationic disordered histone-derived domain (H5). This biosynthetic system allows modular control over charge density, neutral domain length, and stoichiometry, enabling the formation of homogeneous micellar nanoparticles with high protein payload retention under mild, biologically relevant conditions without organic solvents or covalent modification.

Unlike amphiphilic micelle systems, globular proteins in this approach act as both cargo and assembly drivers, providing a direct handle to tune encapsulation. Nanoparticle assembly was characterized with a suite of complementary techniques including DLS, FCS, TEM, and SAXS, enabling detailed analysis of nanoparticle structure, size, and composition. The system exhibits remarkable tolerance to different formulations, consistently producing well-defined particles with low dispersity. This expanded design space arises from the use of structured, charge-regulating proteins and confers exceptional versatility in formulation. Different charge fractions modulate not only nanoparticle size but also physicochemical properties such as core density and cargo loading, which may be critical for tuning nanoparticle stability and performance. Together, these features establish a robust and programmable platform for protein-based nanoparticle engineering with potential application in intracellular protein delivery.

## 1. Introduction

Protein-based biologics have emerged as a key component of modern medicine as they offer precise therapeutic interventions due to their well-defined folded structure. A major bottleneck for protein therapeutics is the delivery of functional proteins into the cytosol in a folded, active form, while avoiding denaturation and degradation. The vast majority of protein drugs currently on the market are limited to extracellular targets due to the challenges with intracellular protein delivery. To address this challenge, various nanoparticle platforms have been developed for protein delivery, including lipid nanoparticles, liposomes, and polymeric micelles.^1^ Due to limitations with these platforms, namely the need to formulate using denaturing solvents, there is increasing interest in materials beyond synthetic nanoparticles to diversify delivery strategies, with a particular focus on protein-based delivery vehicles for targeted and responsive delivery.

Prominent examples of bio-based delivery vehicles include particles based on protein or peptide-driven self-assembly, such as virus-like particles and protein nanocages.^2–5^ In addition to these well-defined protein capsids, protein assemblies driven by hydrophobic interactions have also shown great promise for protein-based drug delivery.^6–14^ For example, phase-separated protein droplets,^6,15,16^ as well as elastin-like polypeptide (ELP)-based micelles and vesicles,^7–14^ have been investigated extensively for the delivery of a range of therapeutic cargo, including proteins. However, the dependence on relatively high concentrations to exceed the critical micelle concentration or harsh processing conditions often limits the biocompatibility and physiological relevance of these hydrophobically driven particles. In contrast, electrostatically driven protein assemblies offer multiple advantages. These include formation under mild, aqueous conditions, inherent responsiveness to salt and pH, and tunability of composition and size via mixing ratios.^17,18^

Electrostatically driven self-assembly has been the predominant approach for the formulation and delivery of nucleic acids.^19,20^ This success has motivated efforts to extend electrostatic assembly strategies to other biomacromolecules, including proteins. Among electrostatically driven assemblies, complex coacervate core micelles (C3Ms) are of particular interest for protein encapsulation and delivery. These self-assembled core-shell nanostructures form when oppositely charged polyions associate, creating a coacervate core stabilized by a neutral hydrophilic corona.^21^ C3Ms, also known as polyelectrolyte or polyion complex micelles (PCM, PIC micelles), were initially developed using simple synthetic polyelectrolytes,^22^ with subsequent in-depth studies on their physical properties, assembly mechanism, and scaling relationships for their design.^21,23–27^ Although globular proteins typically possess a more modest net charge than synthetic polyelectrolytes or nucleic acids, globular proteins can drive assembly as the charged residues are largely localized in discrete patches on the protein surface. Beyond simple systems with two polyions, there are also limited reports of C3Ms with a globular protein as one of the two polyions or as a third globular protein component.^28–31^ Successful intracellular delivery of various biologic cargoes, including proteins and nucleic acids, using C3Ms has been demonstrated.^29,32,33^ But, most studies on protein-containing C3Ms have primarily focused on delivery outcomes rather than fundamental assembly design rules, ultimately limiting the broad applicability of this approach to the full range of potential therapeutic protein cargo.

C3M formation generally requires charge neutrality within the core and a roughly symmetric charge balance between the components, along with an adequately long neutral block to provide the necessary packing constraints.^21,34^ Although this guideline remains valid, the introduction of globular proteins demands careful reconsideration. Globular proteins possess spatially fixed ionizable residues on a folded 3D structure, which influences their complex coacervation behavior. This structural specificity alters the driving forces of C3M assembly and demands an updated framework for understanding and designing micellar protein nanoparticles that incorporate globular proteins. Here, we work to develop these guidelines to expand the utility of micellar protein nanoparticles as robust and programmable protein delivery vehicles.

Herein we report the development of protein-based C3M nanoparticles based on electrostatic self-assembly of multi-domain intrinsically disordered polypeptides (mdIDPs) and globular protein cargo. Guidelines for the design of mdIDPs with a neutral domain that promotes nanoparticle formation and a cationic domain that drives assembly with anionic protein cargo are established. We identify several designs that produce well-defined, monodisperse spherical nanoparticles while incorporating >100 cargo proteins. C3M nanoparticles were formed with a range of globular protein cargo demonstrating the broad utility of this approach for protein formulation and delivery.

## 2. Results and Discussion

### 2.1 Design of biosynthetic multi-domain IDPs for C3Ms

We designed a system where anionic globular proteins could be co-assembled with multi-domain IDPs to form micellar protein nanoparticles (Figure 1). To establish a framework for the self-assembly of globular proteins with mdIDPs, we systematically varied four key parameters: the length of the neutral domain of the mdIDP, the charge of the cationic domain of the mdIDP, the net charge of the globular protein, and the mixing ratios of the globular and the mdIDPs (Figure 1). First, we selected a neutral domain based on a hydrophilic variant of an elastin-like polypeptide (ELP). The modular repetitive architecture of ELPs has well-defined physical properties, including lower critical solution temperature (LCST) behavior. Although ELPs are frequently utilized as phase-separation scaffolds driven by hydrophobic interactions,^9–14,35^ a serine in the “guest” residue position of the pentapeptide repeat, [VPG**S**G]_n_ (ELPS_n_), results in simple coacervation of the ELP above physiological temperatures even for long (n=96) variants (T_t_ > 60 °C).^36^ The largely disordered structure of ELPS_n_ results in an increase in the hydrodynamic volume of the ELP with a power-law dependence on ELP length,^37^ providing a readily tunable platform to engineer the packing parameter of mdIDPs (Supplementary Figure 1, Supplementary Figure 2, Supplementary Figure 3, and Supplementary Figure 4). Moreover, ELPS_n_ lacks ionizable side chains, rendering it neutral regardless of the pH and insensitive to fluctuations in ionic strength or local charge environment. These attributes make ELPS_n_ an ideal inert domain capable of limiting the growth of complex coacervates to create self-assembled nanoparticles.

**Figure 1.**
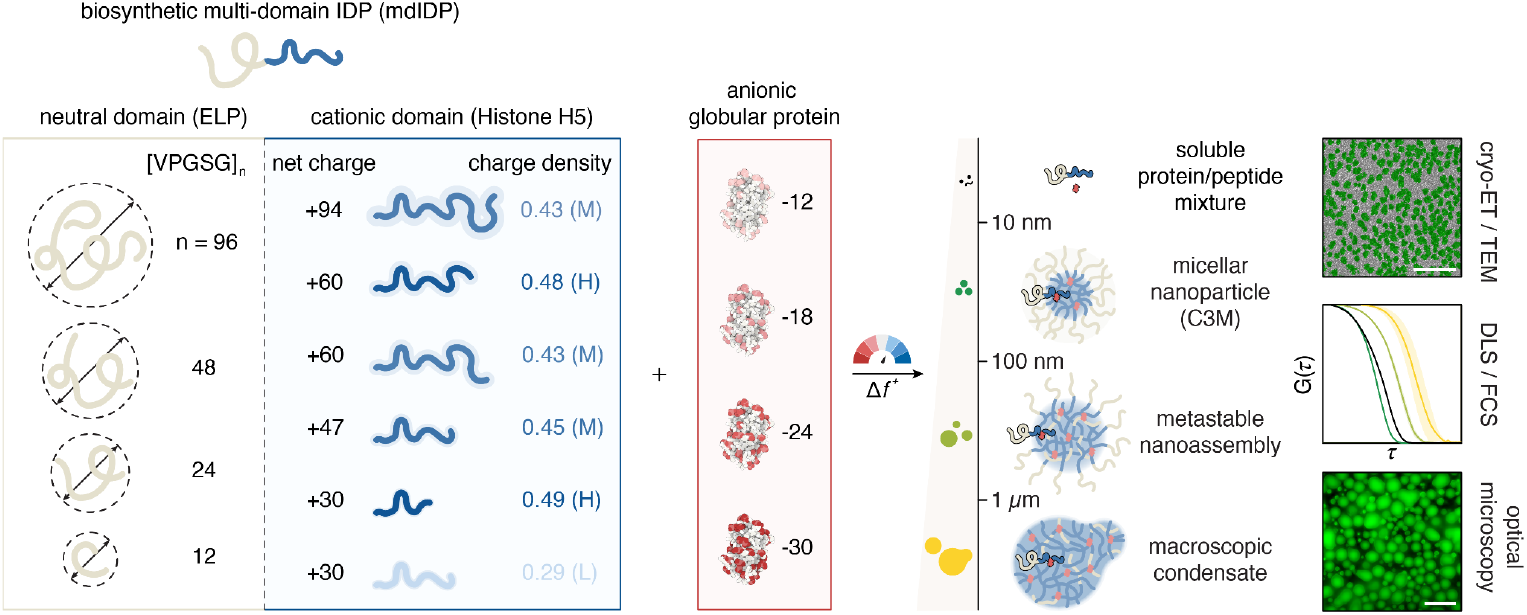
Variations of four parameters determine protein nanoparticle assembly. The length of the neutral domain, the charge of the cationic domain, the net charge of the globular protein, and the mixing ratios of the mdIDP and globular protein were systematically varied. H, M, and L refer to high, medium, and low charge density, respectively. These four parameters collectively influenced the formation of assemblies at different length scales, including micellar protein nanoparticles. Other coacervate-based assemblies were observed, such as macroscopic condensates, metastable nanoassemblies, and soluble protein/peptide mixture. The colors black, green, olive, and yellow correspond to soluble protein/peptide mixture, micellar protein nanoparticle, metastable nanoassembly, and macroscopic condensate. Multiscale validation of the assembly state relied on electron microscopy for nanoscale assemblies, optical microscopy for macroscopic condensates, and DLS and FCS to assess dynamics at all scales.

Second, we engineered a cationic domain, tuning both the net charge and charge density. We started by identifying known IDPs from the DisProt database and selected one with both high net charge and charge density - the disordered region of the nucleosome linker histone H5 (PDB ID: 7XVM).^38^ The wild-type sequence from this region (H5_+47M_) carries a net charge of +47 at pH 7.4, with a net charge per residue (NCPR) of 0.45, and is predicted to be highly disordered with an IUPred2A score of 0.99.^39,40^ By selectively truncating, repeating, and mutating the charged regions of H5_+47M_, 6 different cationic domain variants with net charge ranging from +30 to +94 and charge density ranging from 0.29 to 0.49 were created (Supplementary Figure 3). Fusion of these engineered cationic and neutral domains created a panel of mdIDPs capable of complexing anionic protein cargo.

We next sought to tune the globular protein cargo to evaluate the versatility of the mdIDP platform.^31,41^ Protein cargo with varying net charge, from -12 to -30, were tested for the ability to self-assemble with the mdIDPs and form nanoparticles. A panel of protein cargo, primarily based on engineered variants of superfolder GFP, was used as model biological cargo. Critically, this panel extends beyond those reported in previous studies and includes model cargo beyond GFP.^31,41^

Finally, the mixing ratio between the globular protein and the mdIDP was varied to investigate how the charge balance between the two components impacted the structure, size, and composition of the resulting assemblies. These mixing ratios translated to specific charge fractions (*f* ^+^), a key parameter driving complex coacervation, with a charge fraction of 0.5 corresponding to charge balance between the species which provides the maximum driving force for phase separation. While for linear polyions it is well established that coacervation is most favorable at charge balanced conditions, globular proteins have been shown to form coacervates over an extended range of charge fractions.^31,41,42^ The ability of proteins to form coacervates or C3Ms at charge imbalanced conditions may arise from the fixed geometric arrangement of charges on the folded structure, the presence of charge patches, or the charge regulation of the weak polyampholyte.^21,25,43–48^

With the materials design defined, we then biosynthesized the library of mdIDPs and the panel of model protein cargo. Using a Golden Gate library, the various engineered neutral and cationic domains could be rapidly combined to create fusion gene products encoding for the mdIDPs. The globular proteins and mdIDPs were produced in *E. coli* and purified by affinity chromatography (Supplementary Figure 1). Despite the high charge density of the cationic domain in the mdIDPs, each one of the engineered mdIDPs was successfully produced in *E. coli* with reasonable yields.

### 2.2 Effect of neutral domain length on particle formation

Using this panel of engineered mdIDPs, we first evaluated how the length of the neutral block impacted self-assembly with a single model protein cargo. Initial efforts used GFP-24 as the protein cargo and the wild-type H5 cationic domain with a net charge of +47 (H5_+47M_). Given the potential range of assemblies that could form, from macroscopic condensates to nanoparticles, a suite of characterization tools were needed to differentiate the potential assemblies (Figure 1). Macroscopic condensates were defined as assemblies with diameters greater than a micron, as seen by optical microscopy and denoted by Z-average values of over 1000 nm by dynamic light scattering (DLS) (Supplementary Figure 5). These macroscopic condensates typically exhibit heterogeneous sizes due to dynamic growth of nucleated assemblies. Smaller assemblies, both metastable and equilibrium particles, in the nano-to submicron-scale, were identified by DLS when both the number mean diameter and Z-average were below 1000 nm. To distinguish metastable nanoassemblies and stable micellar nanoparticles, the stability of the hydrodynamic size over time was measured by DLS (Supplementary Figure 6a-c). Finally, nanoscale protein particles were also characterized by complementary techniques, including fluorescence correlation spectroscopy (FCS), electron microscopy (TEM, cryo-ET), and small angle X-ray scattering (SAXS).

Different repeats of ELPS_n_, n = 0, 12, 24, 48, and 96, were tested to identify how the volume of the neutral domain and thus the packing parameter impacted C3M formation. As the length of the neutral domain increased, the macroscopic condensation decreased, shown by both fluorescence microscopy (Figure 2a) and turbidity (Supplementary Figure 5a). From these macroscopic measurements, it appeared that little to no phase separation occurred when *n* ≥ 48. However, nanoscale assemblies are below the detection limit of wide field optical microscopy or turbidity measurement. To observe potential nanoscale particles, samples were characterized by dynamic light scattering (DLS) and fluorescence correlation spectroscopy (FCS) (Figure 2b-c). DLS measurement indicated that ELPS_48_-H5_+47M_ successfully formed nanoscale assemblies with GFP-24 (Figure 2b and Supplementary Figure 5b-d). While the microscopy and turbidity appear to indicate reduced complex coacervation, this apparent decrease simply reflects nanoscale assembly, rather than a genuine reduction in the ion pairing. While coacervation within the core persists, steric crowding of the neutral domain confines the coacervate to a finite nanoscale size, creating a complex coacervate core micelle (C3M).^21,23,49^ The encapsulation of GFP can serve as a qualitative indicator of the underlying thermodynamics of complex coacervation, as the partitioning reflects the net outcome of intermolecular interactions. As evidence of this, we find that the incorporation of GFP-24 in the coacervate domain is consistent for both macroscopic condensates with H5_+47M_ and C3M particles with ELPS_48_-H5_+47M_ (Figure 2c, Supplementary Figure 5e, and Supplementary Figure 7i).

**Figure 2.**
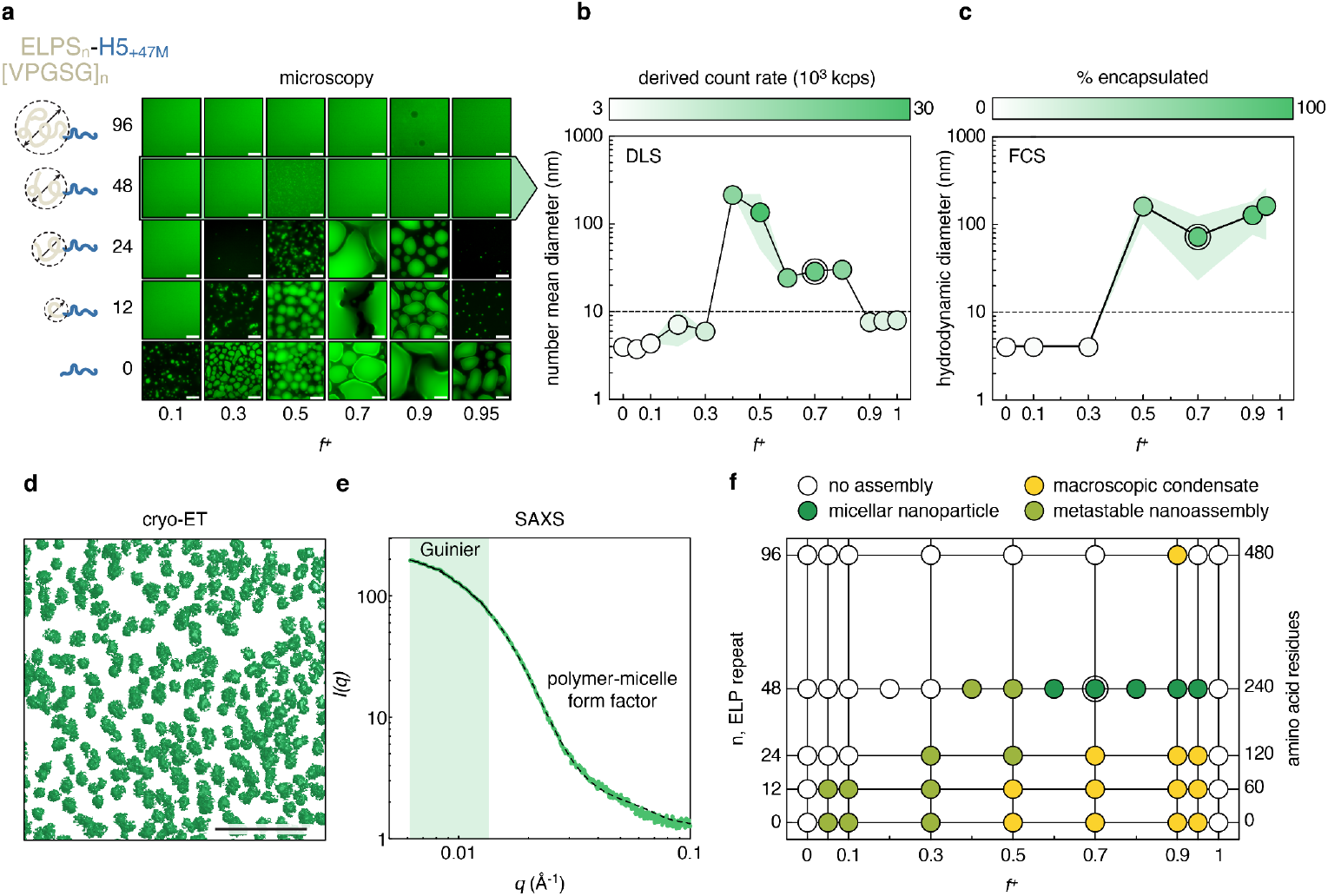
Influence of neutral domain length on the assembly. The length of the neutral domain was varied by changing the number of ELPS_n_ pentapeptide repeats (n), while the cationic domain and anionic globular protein were held constant as H5_+47M_ and GFP-24, respectively. (a) Fluorescence microscopy images of assemblies formed across different neutral domain lengths and charge fractions. Scale bars: 10 µm. (b) Number mean diameter and derived count rate for ELPS_48_-H5_+47M_ assemblies measured by DLS across varying charge fractions. Data points represent the average and green shaded regions represent the standard deviation in the particle size of n = 3 independent experiments, each with 3 technical replicates. (c) Hydrodynamic diameter and fraction of GFP-24 encapsulated in ELPS_48_-H5_+47M_ assemblies as a function of charge fraction, quantified by FCS. Data points represent the average and green shaded regions represent the standard deviation in the particle size of n = 3 independent experiments. (d) Segmented micellar nanoparticles from 3D reconstructed cryo-ET tomogram and (e) SAXS curves of micellar nanoparticles from ELPS_48_-H5_+47M_ and GFP-24 at a mixing ratio of *f*^+^=0.7. Scale bar: 200 nm. (f) Phase diagram as a function of ELP repeat length and positive charge fraction (*f*^+^) illustrating four distinct assemblies based on microscopy, DLS, and FCS analyses. Double-circles in (b), (c), and (f) indicate the condition used for TEM and SAXS analysis in (d) and (e).

Among the nanoscale assemblies, pronounced size heterogeneity and slow growth were observed at mdIDP-deficient charge ratios (*f*^+^ =0.4 - 0.5), whereas mdIDP-excess conditions (*f*^+^ ≥ 0.6) produced monodisperse particles, characteristic of equilibrium nanostructures (Figure 2b and Supplementary Figure 6a-c). The slow, continuous growth of the metastable assemblies when the globular protein was in excess( *f*^+^ < 0.5) reflects insufficient stabilization of the coacervate core due to the limited concentration of the neutral domain. As a result, particles gradually coalesce and grow, albeit slowly producing kinetically arrested nanoscale particles. In contrast, the uniform and equilibrium structures formed under mdIDP-excess conditions are consistent with micellar nanoparticles, in agreement with previous studies on globular protein-based micellar assembly.^31^ These equilibrium nanoparticles were stable for over a month (≥ 43 days; Supplementary Figure 6d-e). Additionally, the micellar nanoparticles remained monodisperse at a concentration down to the detection limit of DLS of 0.15 µM (0.08 µM mdIDP + 0.07 µM GFP-24), without signs of aggregation or disassembly (Supplementary Figure 8). This exceptionally high stability highlights a key advantage of electrostatically driven micellar nanoparticles, which do not rely on micromolar critical micelle concentrations characteristics of amphiphilic micelles.^50^

Micellar nanoparticle formation was corroborated by structural measurements following chemical crosslinking of the core to stabilize particles during analysis. Cryogenic electron tomography (cryo-ET) revealed quasi-spherical particles with a dense core and a diffuse, low-contrast corona typical of polymer micelles.^27^ Assemblies segmented based on the tomographic density yielded fuzzy spherical envelopes characteristic of micelles (Figure 2e and Supplementary Figure 9). The nanoparticles were also characterized by SAXS to probe their global size and internal structure. The scattering profiles were also well described by a polymer-micelle model.^51,52^ This model describes a spherical core surrounded by a polymer corona with a gradual interface and fits the scattered intensity across the measured *q*-range of 0.056 to 1.52 nm^-1^ (Figure 2f and Supplementary Figure 10).

In contrast, as the ELP length was further increased (ELPS_96_) no signs of phase separation were observed (Figure 2a-b). The large volume of the neutral ELPS_96_ domain likely sterically inhibited the associative electrostatic interactions from growing into larger assemblies resulting in solutions of low turbidity (Supplementary Figure 5a), as well as low derived scattering count rate and small Z-average values from DLS analysis (Supplementary Figure 5b-c) across a broad range of charge fractions. However, a single mixing ratio with the mdIDP in excess (*f*^+^ = 0.9) showed an increase in the Z-average and count rate. Although well below the LCST of ELPS_96_, optical microscopy indicated that this scattering was due to simple coacervation as GFP was excluded from the coacervate droplets (Figure 2a). We hypothesize that simple coacervation of the ELPS_96_ was enabled by neutralization of the highly soluble cationic domain by association with GFP-24. These transient electrostatic interactions may facilitate increased local concentration of the high molecular weight ELP and promote the simple coacervation. This behavior was also observed by Horn et al, where simple coacervation of an ELP fusion with an anionic prothymosin-ɑ occurred significantly below the reported critical temperature of the ELP,^53^ when the prothymosin-ɑ domain formed a complex coacervate with a cationic GFP. This transition to simple coacervation, as the ELP solubility decreases, indicates the delicate balance between steric surface stabilization of the desired complex coacervate micelle core and macroscopic condensate formation.

### 2.3. Impact of mixing ratio on assembly size and composition

With confirmation of micellar nanoparticle formation driven by electrostatic interactions between an engineered mdIDP and globular protein cargo, we next sought to understand how the mixing ratio of these two components impacted particle size and formation. We observed micellar nanoparticles with the mdIDP in excess, corresponding to *f*^+^ > 0.5. This observation is consistent with prior reports of C3Ms driven by assembly of a globular protein.^31^ Critically, the use of excess mdIDP allowed for effective encapsulation of the GFP cargo at all mixing ratios that promoted micelle formation (*f*^+^∼ 0.7 - 0.95; Figure 2c). We hypothesize that upon complexation, the globular protein’s effective charge regulates through local modulation of the pK_a_ of surface residues, ultimately allowing for micelle formation at conditions that seem far from charge balance. This broad window for micelle formation has additional advantages as it enables practical tuning of particle size, composition, and density for downstream applications. To understand how the charge balance between the mdIDP and cargo impacted the size and composition of the resulting micellar nanoparticles, we characterized assemblies at varying mixing ratios using a suite of complementary techniques, including DLS, FCS, SAXS, and TEM. A broad range of characterization methods was used as technique-dependent sensitivities and detection biases to particle size or density resulted in differing observable charge fractions.

TEM provided the broadest coverage as the electron-dense coacervate core provided high contrast relative to the soluble proteins and peptides. This allowed for resolution of nanoscale assemblies even near the two-phase boundaries *f*^+^= 0.3 and *f*^+^ = 0.9, where there was no indication of nanoscale assembly formation from DLS due to abundant scattering signal from either the free GFP or free mdIDP components, respectively (Supplementary Figure 9). The assemblies characterized by TEM agreed with the metastable nanoassemblies and micellar nanoparticles observed by DLS as well. Highly spherical and homogenous micellar nanoparticles were visualized at high *f* ^+^ conditions, *f*^+^ = 0.7 and *f*^+^ = 0.9 (Figure 3a,b), consistent with fully stabilized cores with neutral coronas. On the other hand, polydisperse and irregular metastable nanoassemblies were identified at low *f*^+^ regimes conditions, *f* ^+^ = 0.3 and *f*^+^ = 0.5 (Supplementary Figure 9). In addition to TEM, FCS enabled hydrodynamic sizing at conditions that DLS could not, as the single-photon fluorescence detection of FCS isolated the signal from the labeled protein-loaded nanoparticles and ignored excess soluble mdIDP. In the mdIDP-excess regime (*f*^+^ = 0.9 - 0.95), where the nanoparticle was the only fluorescent species (Figure 3d and Supplementary Figure 7). FCS provided more accurate hydrodynamic diameters, with the diameter values at *f*^+^= 0.7 exceeding that of DLS’s ensemble averaging due to the presence of abundant monomers.

**Figure 3.**
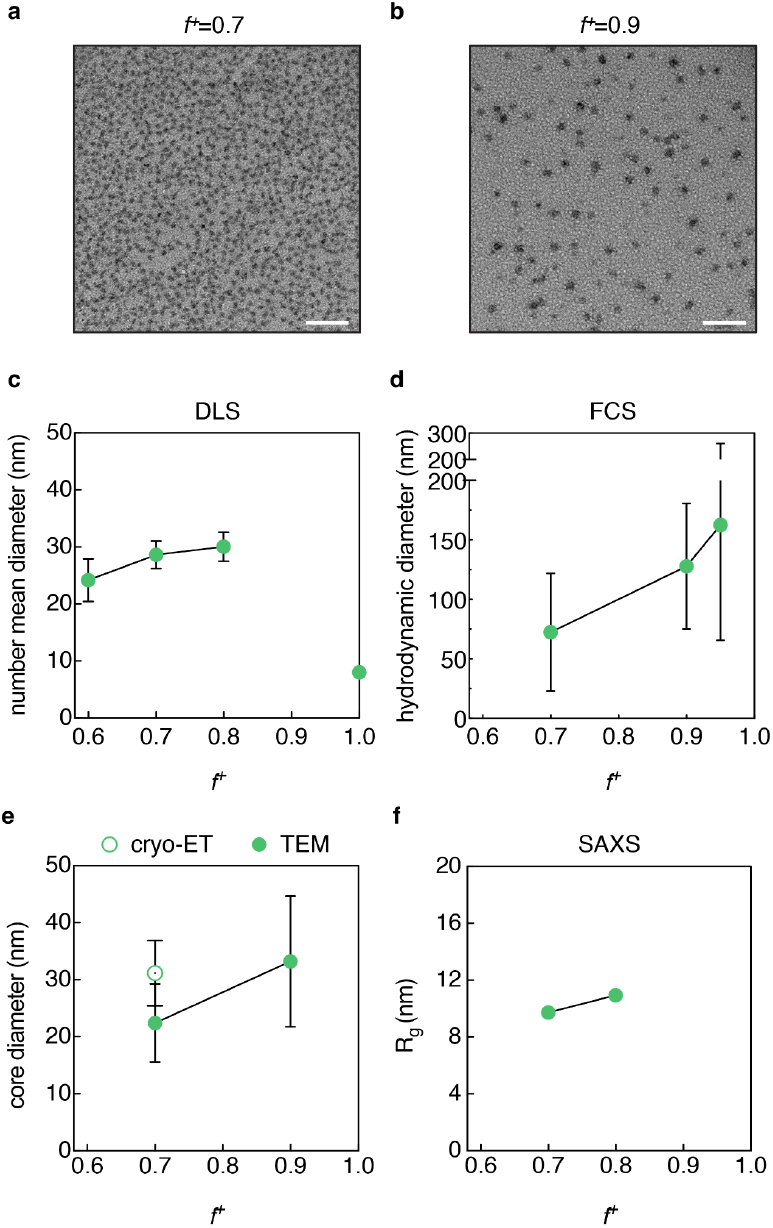
Size characterization of micellar nanoparticles assembled by ELPS_48_-H5_+47M_ and GFP-24. (a) TEM images of micellar nanoparticles assembled at (a) *f*^+^ = 0.7 and (b) *f*^+^ = 0.9. Scale bars: 200 nm. (c) Number mean diameter of micellar nanoparticles from DLS (*f*^+^ = 0.6-0.8) and the free mdIDP (*f*^+^ = 1.0), (d) hydrodynamic diameter from FCS, (e) core diameter from cryo-ET and TEM, and (f) radius of gyration (R_g_) from SAXS at various mixing ratios. For (c-f) data points represent the average and error bars correspond to the standard deviation.

Across all readouts, micellar nanoparticle size increased with the charge fraction, *f*^+^ (Supplementary Figure 11): number mean diameters by DLS (Figure 3c), hydrodynamic diameters by FCS (Figure 3d and Supplementary Figure 7), core diameters by TEM (Figure 3e and Supplementary Figure 9), and radius of gyration (R_g_) by SAXS (Figure 3f and Supplementary Figure 10) all grew with *f*^+^. This growth in particle diameter is likely due to a change in the complexation ratio of GFP and the mdIDP. Under excess mdIDP conditions, the neutral domain supplies sufficient surface coverage to stabilize larger cores; the globular protein’s charge regulation accommodates the off-charge stoichiometry mixing; and the higher mdIDP content promotes hydration and swelling and thus greater core fluidity. Consistent with this depiction, the SAXS form factor fits show the core scattering length density (SLD) decreases with *f*^+^, indicating a lower core density (Supplementary Figure 10). Importantly, this decrease in core density did not compromise cargo protein loading. FCS analysis of the counts per molecule (cpm) rose with *f*^+^ along with the particle size (Supplementary Figure 7j), with FCS estimates indicating ∼200-400 GFP-24 molecules per nanoparticle. The charge fraction dependent control of size, composition, and fluidity provides a practical handle to optimize particle performance, made possible here as the globular protein serves as the assembly driver.^1,54,55^

### 2.4. Effects of cationic domain charge

Following successful micellar nanoparticle formation with ELPS_48_-H5_+47M_ and GFP-24, we next sought to investigate how the net charge and charge density of the cationic domain impacted particle formation. Five additional cationic domains were genetically fused with the ELPS_48_ neutral domain. The electrostatic-driven self-assembly of each one of these mdIDPs with GFP-24 was then evaluated (Figure 4a). Four of the five additional variants, ELPS_48_-H5_+30H_, ELPS_48_-H5_+60M_, ELPS_48_-H5_+60H_, and ELPS_48_-H5_+94M_, exhibited similar behavior to the original wild-type H5 variant (ELPS_48_-H5_+47M_) forming metastable nanoassemblies and micellar protein nanoparticles (Figure 4b). It was notable that the H5_+30H_ variant was able to assemble into nanoparticles, while the H5_+30L_ variant did not despite the two domains having identical net charge. The difference in behavior between the H5_+30H_ and H5_+30L_ domains was more evident from analysis of variants with different lengths of the neutral domain (n = 0, 12, 24, or 48). As shown by optical microscopy, macroscopic condensation was significantly diminished for the ELPS_12_-H5_+30L_ fusion, while condensate formation was maintained for ELPS_24_-H5_+30H_ (Figure 4c-d and Supplementary Figure 12a-b). As the ELP length was increased to n = 48, the count rates from DLS analysis also showed a significant difference between the two +30 variants. Nanoscale assemblies were only detected for H5_+30H_ with the ELPS_48_ neutral domain. For variants with lower overall charge (+30), it seems that, strong complexation mediated by clustered cationic residues in H5_+30H_ is critical for stable formation of ion pairs that lead to macroscopic condensation or micellar nanoparticle formation.^56^

**Figure 4.**
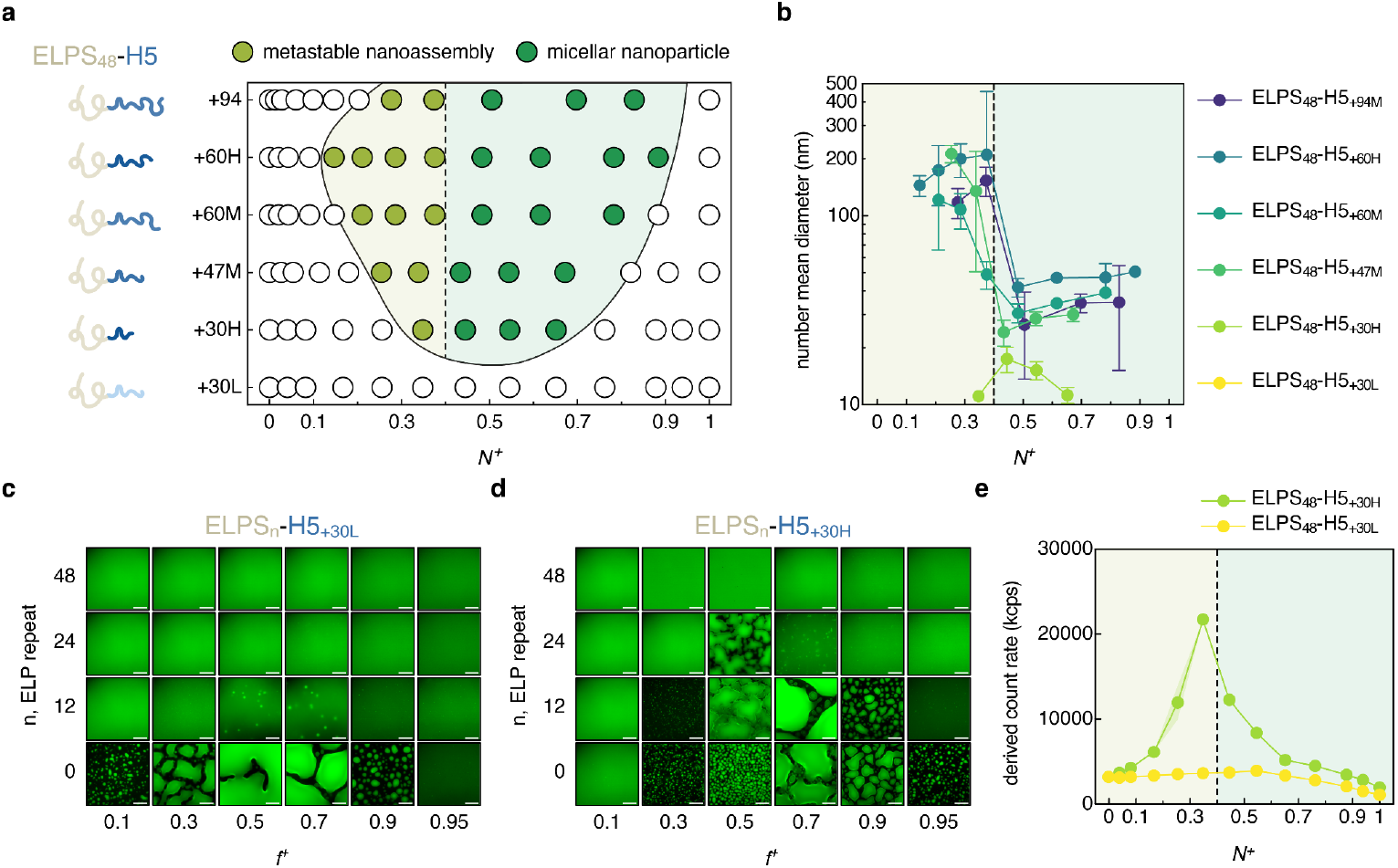
Various assemblies observed from mdIDPs with engineered cationic domains. (a) Phase diagram illustrating metastable nanoassembly and micellar nanoparticle assembly regimes for different cationic domains as a function of stoichiometric mixing ratios, determined by DLS. (b) Number mean diameter of metastable nanoassembly and micellar nanoparticle as measured by DLS. Data points represent the average of 3 independent experiments and the error bars correspond to the standard deviation of these measurements. The neutral domain and anionic globular protein were fixed as ELPS_48_ and GFP-24, respectively in (a) and (b). Dotted lines in (a), (b) and (e) denote *N*^+^ = 0.4, the threshold at which a reduction in assembly size and transition from metastable nanoassembly (shaded in olive color) to micellar nanoparticles (shaded in green color) was observed. (c) Fluorescence microscopy images of assemblies formed by ELPS_n_-H5_+30L_ and ELPS_n_-H5_+30H_ with GFP-24. Scale bars: 20 µm (e) DLS-derived count rates of ELPS_48_-H5_+30L_ and ELPS_48_-H5_+30H_ complexed with GFP-24. Data points represent the average of 3 independent experiments and the shaded regions correspond to the stand deviation of these measurements.

While the difference in the charge densities between H5_+30L_ and H5_+30H_ was a determinant for crossing the threshold for coacervation with ELPS_48_, cationic domains with higher net charge enabled the assembly of more stable and homogenous nanoparticles at a broader range of mixing ratios. ELPS_48_-H5_+60H_ assembled into nanoparticles of the lowest dispersity when compared to a variant with identical net charge, ELPS_48_-H5_+60M_, and even when compared to one with higher net charge, ELPS_48_-H5_+94M_ (Supplementary Figure 13a). Furthermore, ELPS_48_-H5_+60H_ had the broadest assembly regime, with both metastable nanoassembly and micellar nanoparticle micelle formation occurring over the widest range of mixing ratios.

Finally, it was noticeable that the transition from metastable to equilibrium nanoparticle assemblies consistently occurred around a molar fraction of the mdIDP (*N*^+^) of 0.4 for all five mdIDP variants that formed assemblies. This deviates from the conventional expectation for synthetic polymer systems, where optimal nanoparticle formation is observed at charge neutral conditions (*f*^+^ = 0.5) (Supplementary Figure 13d).^27,43^ This constant molar ratio requirement for nanoparticle formation spanned mdIDPs with over a 3-fold difference in the net charge of the cationic domain, resulting in micelle formation at *f*^+^ ranging from 0.5 to 0.95. With the optimal molar mixing ratio constant (constant *N*^+^) regardless of the net charge of the cationic domain, the nanoparticle core has an increased cationic charge as the net charge on the mdIDP increases (increasing *f*^+^ at this optimal molar ratio). To balance this increased cationic charge the GFP-24 should decrease its charge by undergoing charge regulation.^46,47,47,48^ Charge regulation is commonly observed for globular proteins as they are weak polyampholytes, ultimately broadening the assembly regime to conditions with excess mdIDP. This ability to regulate charge makes globular proteins excellent candidates to drive micellar nanoparticle assembly. On the other hand, stronger interaction, coming from higher net charge of the cationic domain or the anionic protein, is not always the key for more favorable or homogenous nanoparticle formation. Optimal *f*^+^ for ELPS_48_-H5_+94M_ was stretched to 0.8 and 0.95 range, for *N*^+^ ∼ 0.4 constraint for GFP-24 (Supplementary Figure 13d). Although charge regulation of GFP-24 was able to stretch the phase separation to a high *f* ^+^ condition, the broken symmetry and charge imbalance environment failed to provide a favorable condition for complex coacervation, leading to less optimal nanoparticle formation than ELPS_48_-H5_+60H_. Counterintuitively, given the central role of charge stoichiometry in driving micellar nanoparticle assembly, we found that the assembly is largely independent of the absolute protein concentration. Conditions previously falling outside of the micelle-forming-regime, such as 17.9 µM GFP at *f*^+^ = 0.9 with ELPS_48_-H5_+47M_ by DLS (Figure 2b) - could be readily shifted into the nanoparticle-forming regime through simple adjustment of the mixing ratio (Supplementary Figure 8c). This imparts an excellent advantage in formulating nanoparticles for protein therapeutics with a range of electrostatic properties by tuning the charge ratio of mdIDPs.^54,55^

### 2.5. Nanoparticles form with range of globular protein cargo

With an understanding of both the critical neutral domain length and cationic domain charge, we next sought to gauge how the globular protein cargo impacted nanoparticle assembly. Initially, the net charge was varied between -12 to -30 on a common GFP scaffold and nanoparticle formation with ELPS_48_-H5_+47M_ was evaluated (Figure 5a). All four GFP variants tested successfully formed micellar nanoparticles. Nanoparticle formation was observed consistently around *f*^+^ = 0.7 across GFP variants, denoted by low polydispersity values (Figure 5a and Supplementary Figure 14a). The same behavior was observed for another mdIDP, ELPS_48_-H5_+60H_, with a consistent *f*^+^ for nanoparticle formation, though it was shifted to a slightly higher positive charge fraction (Figure 5b). This dependence on the globular protein was in contrast to the observed behavior for varying cationic domains, with globular proteins showing a preferred charge ratio while varying mdIDPs had a preferred molar ratio (*N* ^+^; Figure 4a and Supplementary Figure 13d). This suggests that for a fixed mdIDP nanoparticle formation occurs at a particular charge fraction (*f*^+^) with a range of different globular proteins. As GFP behaves as a weak polyampholyte, it is possible that the protein modulates the surface charge such that complexation with the cationic mdIDP is feasible at a broad range of charge fractions. The ability of both the globular protein cargo and the mdIDP to undergo charge regulation expands the conditions for electrostatic assembly of protein nanoparticles potentially enabling a broad range of applications.

**Figure 5.**
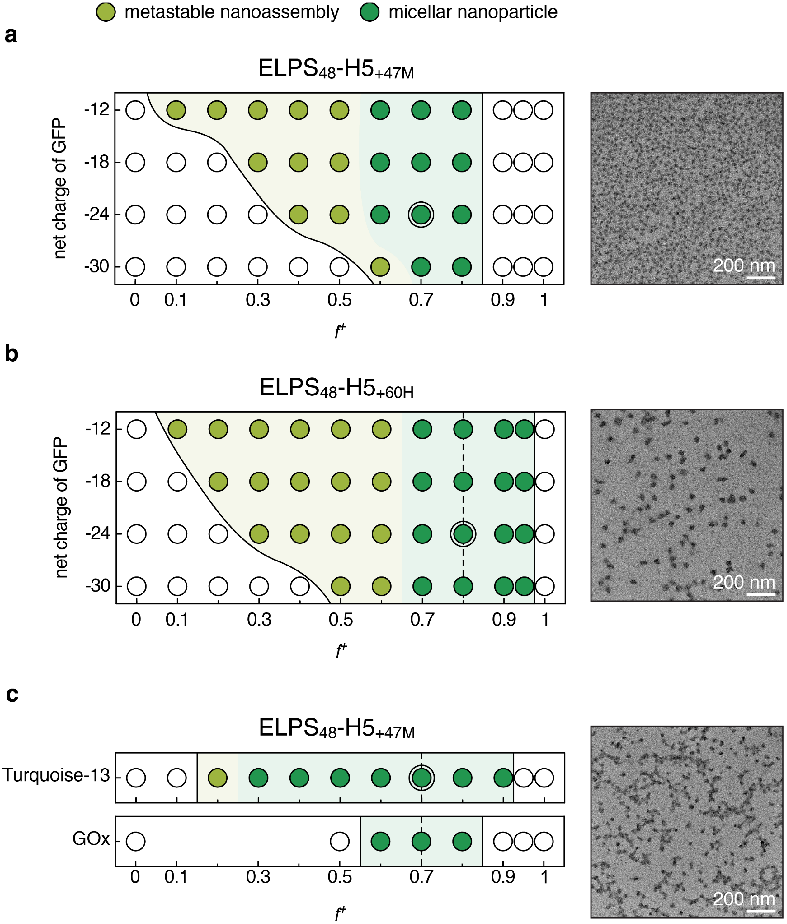
Micellar nanoparticle formation with various globular protein cargos. Phase diagram illustrating metastable nanoassembly and micellar nanoparticle assembly regimes and representative TEM image for (a) ELPS_48_-H5_+47M_ and (b) ELPS_48_-H5_+60H_ with GFPs of various net charge. (c) Phase diagram illustrating metastable nanoassembly and micellar nanoparticle assembly regimes and representative TEM image for ELPS_48_-H5_+47M_ with Turquoise-13 and GOx. Dotted lines denote charge fraction where the lowest polydispersity was observed, *f*^+^= 0.7 and *f*^+^ = 0.8 for ELPS_48_-H5_+47M_ and ELPS_48_-H5_+60H_, respectively. Representative TEM images were acquired from the double-circled points on the phase diagrams. Additional TEM images of micellar nanoparticles from various globular protein cargoes can be found in Supplementary Figure 14.

Beyond the model GFP cargo with varying charge, additional globular proteins could similarly drive nanoparticle formation. Both the enzyme glucose oxidase (GOx, -29) and an mTurquoise2 variant (Turquoise-13) also formed well-defined protein nanoparticles as demonstrated by both DLS and TEM (Figure 5c and Supplementary Figure 14e-g). This consistent formation of protein nanoparticles regardless of the globular protein cargo demonstrates the generalizability of this strategy to proteins with diverse sizes, folds, and surface chemistries. Here we present the broad applicability and scope of the mdIDP protein particle platform as it supports cargo-agnostic encapsulation across a wide charge range, preserving functional proteins, and enabling tunable formulations for downstream applications from intracellular protein delivery to enzymatic nanoreactors.

## 3. Conclusion

We investigated the assembly of protein nanoparticles driven by electrostatic interactions between engineered multi-domain IDPs and protein cargo. Dependence of the nanoparticle assembly on four key parameters - (1) neutral domain length, (2) cationic domain charge, (3) globular protein charge, and (4) the mixing ratio of mdIDP and protein - was evaluated. This was enabled by the simple, tunable design of biosynthetic mdIDPs, where modular designs were genetically encoded and produced *E. coli*. We found that the length of the neutral domain was a key determinant of the final assembly state - with shorter neutral domains promoting macroscopic condensate formation and intermediate length neutral domains facilitating nanoparticle formation. If too long, the neutral domain could completely inhibit electrostatic self-assembly. Tuning of the net charge and charge density of the cationic domain revealed that the charge density of this domain was critical for the formation of stable, monodisperse nanoparticles. Near the charge threshold for particle formation, decreasing the charge density could completely suppress micellar nanoparticle formation. Critically, the design criteria for assembly with mdIDPs was generalizable across a range of model globular protein cargo with varying net charge and molecular weight. We demonstrated that mdIDPs successfully self-assembled into uniform micellar nanoparticles with nearly 100% of the globular protein incorporated into the nanoparticle. Depending on the mdIDP sequence and the mixing ratio with the protein cargo, the composition and size of the assembled protein particles could be varied. To the best of our knowledge, this behavior has not been observed in synthetic polymer C3M systems. With the mdIDP in excess, we found that the particles increased in size and incorporated more protein cargo. This likely occurs due to charge regulation of the globular protein, which allows for a broader range of formulations that promote self-assembly.

The multi-domain IDP platform established here provides a modular framework that allows for the straightforward integration of sequence- or structure-specific function into micellar protein nanoparticles. The mdIDP platform provides engineering opportunities as the micellar nanoparticle core is tunable with the ability to modulate physical properties such as the core hydration, viscoelasticity, protein loading, and charge density. These parameters can be adjusted to influence not only the micelle stability and cargo retention, but also to maximize cellular uptake and cargo release. Finally, as the nanoparticle assembly is largely independent of the protein cargo, the mdIDP platform enables the formulation and delivery of diverse therapeutic globular protein cargoes, guided by the design rules established here. Looking forward, we plan to leverage this platform for therapeutic protein delivery, incorporating functional domains that include receptor-binding proteins to mediate targeted cellular interactions and endosomolytic motifs to enable controlled endosomal escape.

## Acknowledgements

This work was supported by the National Institutes of Health under award number NIH-NIGMS-R35GM138378, US Department of Energy, Office of Basic Energy Sciences, Grant DE-SC0008772, and the US Department of Defense, Army Research Office, W911NF-22-2-0111. SAXS experiments used resources of the Center for Functional Nanomaterials and National Synchrotron Light Source II, supported by the U.S. DOE Office of Science Facilities at Brookhaven National Laboratory under Contract No. DESC0012704. This study used the resources of the CUNY advanced Science Research Center Imaging Suite for FCS. Cryo-ET experiments were supported by the Simons Electron Microscopy Center and the National Resource for Automated Molecular Microscopy located at the New York Structural Biology Center, supported by grants from the Simons Foundation (SF349247) and the NIH National Institute of General Medical Sciences (GM103310).

