## Supplementary Information for "Electrostatic Self-Assembly of Protein Nanoparticles from Intrinsically Disordered Polypeptide and Globular Protein Pairs"

### Methods

#### Cloning

The DNA sequences of GFP-12, GFP-18, GFP-24, and GFP-30 in a pETDuet vector were previously constructed.[1,2] DNA sequences of mdIDPs were constructed in a pET24a(+) vector backbone. The gene for an elastin-like polypeptide (ELP) with serine in the guest residue position ($ELPS_{96}$) in a pET25b(+) vector was purchased from Addgene (Plasmid #68393). $H5_{+30L}$ and $H5_{+47M}$ variants of the H5 gene in pET24a(+) vector were purchased from TWIST Bioscience. $ELPS_{12}$, $ELPS_{24}$, $ELPS_{48}$, and $ELPS_{96}$ were PCR amplified from the Addgene template ($ELPS_{96}$) and assembled with the $H5_{+30L}$ and $H5_{+47M}$ in the pET24a(+) vector through Hi-Fi assembly. Individual domains were subsequently cloned into donor vectors, as shown in Supplementary Figure 4 through Hi-Fi assembly, for Golden Gate assembly into various combinations of H-ELP mdIDPs. Iterative cycles of BsaI restriction digestion at 37 °C and ligation at 16 °C were performed to ensure effective one-pot Golden Gate assembly for the remaining constructs.

#### Protein expression of globular proteins

GFP variants and mTurquoise2 were expressed in NiCo21(DE3) *E. coli* cells in 1 L cultures of LB media. LB media was supplemented with 100 µg of ampicillin. Cultures were incubated at 37 ºC, with shaking at 175 rpm. Cultures were grown to $OD_{600}$ of ~0.9 and were subsequently induced by addition of 1 mL of 1 M isopropyl β-D-1-thiogalactopyranoside (IPTG). The cultures were incubated with shaking at 25 ºC for 16 h after induction.

#### Protein expression of mdIDPs

mdIDPs were expressed in BL21-A1 *E. coli* cells in 1 L cultures of LB media. LB media was supplemented with 100 µg of kanamycin. Cultures were incubated at 37 ºC, with shaking at 175 rpm. Cultures were grown to $OD_{600}$ of ~0.6 and were subsequently induced by addition of 1 mL of 1 M isopropyl β-D-1-thiogalactopyranoside (IPTG) and 10 mL of 20% L-arabinose. The cultures were incubated with shaking at 25 ºC for 16 h after induction.

#### Cell lysate preparation for globular proteins

Cells from 2 L of culture were harvested by centrifugation at 3500 g for 15 min, and cell pellets were resuspended in 30 mL of lysis buffer (50 mM $NaH_2PO_4$, 300 mM NaCl, pH 8.0) and frozen at -80 ºC overnight. The thawed cells were lysed by sonication with a microtip (1/2 in, 2 s on, 4 s off, total 10 min sonication time with 40% amplitude) and cell debris was removed by centrifugation at 11,000 g for 30 min. The separated supernatant was filtered with a polyethersulfone (PES) syringe filter with a pore size of 0.22 µm to remove residual insoluble particles.

#### Cell lysate preparation for mdIDPs

Cells from 2 L of culture were harvested by centrifugation at 3500 g for 15 min, and cell pellets were resuspended in 30 mL of lysis buffer (50 mM $NaH_2PO_4$, 1 M NaCl, pH 8.0) and frozen at -80 ºC overnight. The thawed cells were supplemented with 600 µL of a protease inhibitor cocktail solution (P8849, Sigma Aldrich), 30 µL of a 10 mg/mL DNase I solution, 30 µL of a 10 mg/mL RNase A solution, 300 µL of a 1 M $MgCl_2$ solution, 300 µL of a 1 M $CaCl_2$ solution, and 300 µL of Tween 20. The supplemented cells were lysed by sonication with a microtip (1/2 in, 2 s on, 4 s off, total 10 min sonication time with 40% amplitude). Cell lysates were heat-treated by incubation in a 90 °C water bath for 10 min, followed by rapid cooling in an ice bath. This heat-cool cycle was repeated twice. Insoluble cell debris was then removed by centrifugation at 11,000 g for 30 min. The clarified supernatant was filtered with a 0.22 μm PES filter to remove residual insoluble particles.

**Protein purification**

Protein purification was performed on an ÄKTA FPLC system (Cytiva) using a 25 mL HisPur™ Ni-NTA resin column (Thermo Scientific). Solution A consisted of lysis buffer without imidazole, and Solution B was the lysis buffer containing 250 mM imidazole in the same buffer base. The column was equilibrated with 2 column volumes (CV) of Solution A. The clarified lysate was then loaded onto the column at a constant flow rate. After sample application, the column was washed with 6 CV of buffer containing 40 mM imidazole (achieved by mixing 16% of Solution B with 84% of Solution A) to remove nonspecifically bound proteins. Elution was performed using 2 CV of 100% Solution B. Six fractions were collected during the elution, each with a volume of 8.3 mL. The fractions were analyzed by SDS-PAGE on a 4–12% Bis-Tris gradient gel to assess protein purity and molecular weight.

**Protein preparation**

The collected FPLC eluates were dialyzed against 4 L of 10 mM Tris buffer (pH 7.4) using a regenerated cellulose dialysis membrane with a 3.5 kDa molecular weight cut-off (MWCO). At least seven buffer changes were performed after a minimum 3 h interval to ensure complete buffer exchange. Buffer exchanged proteins were filtered with a 0.22 μm PES syringe filter to remove aggregates. Filtered proteins were concentrated by ultrafiltration using Amicon centrifugal filter units to a final protein concentration of 150 μM. mdIDPs without neutral domains and with $ELPS_{12}$ were concentrated using 3 kDa MWCO filters, while 10 kDa MWCO filters were used for the other $ELPS_n$ fusion proteins. The concentration of globular proteins and mdIDPs was determined by Bradford Assay and Quant-iT Protein Assay (Thermo), respectively, as described by the manufacturer. mdIDPs were flash frozen with $LN_2$ and kept at -80 ºC for storage. Thawed proteins were re-filtered with a 0.22 μm PES filter and the concentration was re-quantified before use. Glucose oxidase was purchased from Sigma Aldrich (G7141).

**LC-MS**

Sample solutions were made by diluting the protein stock solution to 0.1 mg/mL in Milli-Q purified water. Reversed-phase chromatography was performed using an Agilent 1260 HPLC system equipped with a 1×75 mm InfinityLab Poroshell 300Extend-C18 column (671750-902, Agilent Technologies). Solvent A and B were composed of 0.1% formic acid in Milli-Q water and acetonitrile, respectively. The flow rate was 0.4 mL/min and the column temperature was set at 25 ℃. The column was equilibrated with 3% B for 10 min prior to sample injection. A sample volume of 5 μL was injected onto the column. The gradient elution profile was from 3 to 90% B for 4 min followed by washing with 97% B for 1 min. Full mass data was acquired with a range of 100-3000 m/z and an acquisition rate of 1 spectrum/s on an Agilent 6230 LC/TOF system. The instrument was operated in positive mode for intact protein analysis. Mass spectra were deconvoluted using the Agilent MassHunter BioConfirm software with the built-in Maximum Entropy algorithm.

**Complex coacervation induction and protein assembly**

The concentration of both the globular protein and mdIDPs were adjusted to 100 μM unless otherwise specified. Prior to mixing, both solutions were equilibrated to 25 °C. To achieve different mixing ratios, varying volumes of the protein and mdIDP solutions were combined in a PCR tube while maintaining a constant total macromolecule concentration, unless noted otherwise. The mdIDP was always added to the globular protein solution to ensure consistent mixing, followed by gentle pipetting. The mixture was incubated at 25 °C for 1 h before subsequent characterization. Total sample volumes varied depending on the requirements of the downstream analytical method(s).

**Turbidity assay**

Samples were mixed in a half-area 96-well plate at various mixing ratios of GFP and mdIDPs. The total sample volume was maintained at 50 µL per well. Each mixing condition was prepared in triplicate and absorbance at 600 nm was measured using a plate reader (Tecan Infinite M200 Pro). Turbidity was calculated using the equation: Turbidity (%) $= 100 - 10^{2-A}$, where A is the absorbance at 600 nm with pathlength correction.

**Fluorescence microscopy**

Microscopy images were taken using a Nikon Ti Eclipse fluorescence microscope. Samples (20 µL) were prepared at 100 µM total macromolecular concentration with varied charge fractions in a 384-well glass bottom plate (Cellvis P384-1.5H-N). All images were taken using a 100× Plan-Apochromat NA 1.40, objective lens, with additional 1.5× optical magnification. A 488 nm laser was used at 1% power for excitation, and fluorescence was detected using a 470/40 excitation and 525/50 emission filter set.

**Encapsulation assay**

Encapsulation assays were done as previously reported.[3] Solutions of GFP(-24) (50 µM) and $ELPS_0$-$H5_{+47M}$ (50 µM) were mixed in a 96-well round bottom plate at various mixing ratios as above. The total sample volume was maintained at 100 µL per well. After 1 h of incubation at 25 °C, the samples were centrifuged at 3500 g for 30 min to separate the dense and dilute phases. 80 µL of the dilute phase was removed for measurement and the remaining dilute phase (approximately 20 µL) was discarded. Then, 100 µL of a 2 M KBr solution was added to the dense phase and gently pipetted to redisperse the coacervate. After resuspension, 80 µL of this solution was removed for analysis. The concentration of GFP-24 in the dilute phase and the re-dissolved dense phase was analyzed through fluorescence measurements using a plate reader with $\lambda_{ex}$ = 470 nm and $\lambda_{em}$ = 520 nm (Tecan infinite M200 Pro).

**DLS**

For dynamic light scattering analysis, 50 µL of each sample (prepared in triplicate) was transferred to a low-volume disposable cuvette (Malvern ZEN0040). Measurements were performed using a Malvern Zetasizer Nano ZS at 25 °C in 173° backscatter detection mode. The measurement position was set to 3 mm, with 3 runs per sample, each consisting of 3 acquisitions of 10 seconds. The viscosity of the medium was set to 0.8872 cP. The size distribution of multimodal particle populations was determined by fitting the autocorrelation curves using the instrument's distribution analysis.

**FCS**

Fluorescence Correlation Spectroscopy (FCS) was conducted using a Leica TCS-SP8 STED 3X inverted confocal microscope equipped with a 63× 1.20 NA water-immersion objective. Samples were excited with a 488 nm line from a white light laser pulsed at 40 MHz. Emission was collected through an adjustable 70 µm pinhole and detected between 520–540 nm using a HyD detector connected to a TCSPC module. Data acquisition and autocorrelation analyses were performed in SymPhoTime 64 (PicoQuant). The focal volume was calibrated using the free GFP-24 diffusion ($D = 122$ µm$^2$ s$^{-1}$ at 25 °C).

For each sample, triplicate measurements (300 s each) were acquired across three different days. Autocorrelation curves $G(t)$ were fit to diffusion models (1–3 components) including a triplet-state term[4]:

$$G(t) = 1 + \frac{1}{<N>}\left(1 + \frac{F_{trip}}{1-F_{trip}} \cdot e^{-t/T_{trip}}\right) \cdot \sum_{i=1}^{n} \frac{F_i}{(1+(t/\tau_{diff,i}))\cdot\sqrt{1+(\omega_{xy}/\omega_z)^2\cdot(t/\tau_{diff,i})}}$$

Where $< N >$ is the average number of fluorescent molecules $N$ in the focal volume. $F_{trip}$ is the fraction of molecules in the triplet state, $T_{trip}$ is the average time a molecule resides in the triplet state. $F_i$ and $\tau_{Diff,i}$ are the fraction and diffusion time of species $i$ out of $n$ components, respectively. $\omega_{xy}$ and $\omega_z$ are the equatorial and axial radii of the detection volume, respectively. For nanoparticles, the autocorrelation curves were fit with models with $n$ = 2–3 components, constraining one component's diffusion time to that of free GFP. Hydrodynamic radii were calculated via the Stokes–Einstein equation.

To estimate the number of GFP molecules per micelle, two methods were used: (1) comparing particle and free GFP number concentrations, assuming 100% encapsulation; and (2) comparing molecular brightness (counts per molecule, cpm). To avoid quenching or Förster resonance energy transfer effects between GFPs, 40 mol% of dark GFP-24 was added. The cpm was calculated by normalizing the total photon rate (counts per second, cps) by the particle number $N$.

**Glutaraldehyde crosslinking**

Prepared assemblies were cross-linked by incubation with glutaraldehyde (340855, Sigma Aldrich) at a final concentration of 0.01 wt% for 30 min at room temperature for TEM, cryo-ET, and FCS analysis.

**TEM**

Samples (5 µL) of glutaraldehyde cross-linked assemblies were applied to a 300-mesh carbon-coated copper grid (Ted Pella 01843-F). Excess liquid was wicked away using filter paper, and the grid was stained with 5 µL of 2% uranyl acetate for 30 s. After removing the stain by wicking, the grid was washed twice with 5 µL of distilled water, wicking between each wash. Grids were air-dried at room temperature and stored at room temperature until imaging. TEM imaging was performed using an FEI Talos F200X. The area of assemblies were analyzed using FIJI. A Gaussian blur filter was applied with radius values of 20 nm for assemblies formed at $f^+ = 0.3$, 5 nm for $f^+ = 0.5$ and 0.9, and 3 nm $f^+ = 0.7$. Thresholding was performed using the Max Entropy method. The analyze particle tool was used for particle quantification. Particle size thresholds were set to 100 $nm^2$-infinity at $f^+ = 0.5$, 0.7, and 0.9, and 500 $nm^2$-infinity for $f^+ = 0.3$.

**Cryo-ET**

A solution of glutaraldehyde cross-linked protein nanoparticle (3.5 µL) was applied onto AuUltrafoil R2/1, 200 mesh grids. The grids were blotted with a vitrobot system (Thermo Fisher Scientific) with blot times ranging from 3-4.5 s at 4 °C with 100% humidity and plunge frozen into liquid ethane. The samples were visualized with a Titan Krios electron microscope (Thermo Fisher Scientific) equipped with a field emission gun, a GIF Quantum LS postcolumn energy filter (Gatan) and a K3 summit electron detector (Gatan). The electron microscope was operated at 300 kV in nanoprobe mode at a magnification of 42,000x (pixel size of 2.123 Å at the specimen level). Cryo-ET tilt-series were collected with a dose symmetric scheme with a tilt-range of -52 to +52° at a target defocus of -4 µm and 3° increments in SerialEM for a total dose of 145 e/$Å^2$.[5,6]

The raw tilt movies were motion and CTF (contrast transfer function)-corrected in Warp 1.09.[7] The tilt-series stack was exported from Warp, where the tilt-series were aligned with the AreTomo software package.[8] The aligned tilt-series were reconstructed with weighted-back projection (WBP) and CTF-deconvolved with Warp. To enhance the contrast of the tomograms for visualization, the WBP CTF-deconvolved tomograms were imported into IsoNet,[9] where missing-wedge correction was performed. The missing-wedge corrected tomograms were imported into ORSDragonfly where the nanoparticles in the tomograms were manually segmented. Segmented objects were converted into multi-ROI sets using 6-connectivity criteria. The volume of each object was calculated as the total number of constituent voxels, with a voxel size of 1.4 nm per side. Objects with volumes smaller than 1000 $nm^3$-equivalent to spheres with radii under 6.2 nm-or with an aspect ratio greater than 3.5 were excluded from further analysis.

**SAXS**

The measurements were performed at Complex Materials Scattering (CMS, 11-BM) beamlines at the National Synchrotron Light Source II (NSLS-II) in Brookhaven National Laboratory with a photon energy of 13.5 keV and a sample-to-detector distance of 5.03 m. The horizontal and vertical beam size was 200×200 μm. Scattering signals were collected using a Pilatus 2M detector, with a pixel size of 172× 172 μm. For sample preparation, 60 μL of the sample solution was loaded into a glass capillary (Boron-Rich 15-BG-1.5 mm; Charles Supper Company). The capillary was sealed with wax to prevent evaporation and measured with an exposure time of 10 s.

Two-dimensional (2D) scattering patterns were collected using area detectors positioned downstream of the sample. These images were azimuthally integrated to produce one-dimensional (1D) scattering intensity profiles, I(q), as a function of the scattering vector $q = \frac{4\pi}{\lambda} sin(\frac{\theta}{2})$, where λ is the incident X-ray wavelength and θ is the total scattering angle. To isolate the sample scattering signal, the background scattering from the corresponding buffer solution, measured under identical conditions, was subtracted from the raw data prior to further analysis. The resulting 1D profiles covered a $q$-range of approximately 0.04 to 1 $nm^{-1}$, with a resolution of 0.002 $nm^{-1}$. The radius of gyration was determined by Guinier analysis in the low q regime, with the fitting range restricted to q values satisfying $q_{max}R_g < 1.3$ to ensure the validity of the Guinier approximation. The 1D scattering profile was fit with a polymer micelle form factor model using the SasView 6 software.[10,11] This model describes a micelle composed of a spherical core with uniform scattering length density, surrounded by Gaussian polymer chains attached to the core, forming a corona. The model provides the form factor, $P(q)$, which describes the scattering intensity, as a function of the scattering vector $q$, and is calculated using the following equations:

$$P(q) = N^2\beta_s^2\Phi(qr)^2 + N\beta_c^2P_c(q) + 2N^2\beta_s\beta_cS_{sc}(q) + N(N-1)\beta_c^2S_{cc}(q)$$

With the contrast amplitude (β) of the core and corona as follows:

$$\beta_s = V_{core}(\rho_{core} - \rho_{solvent}), \beta_c = V_{corona}(\rho_{corona} - \rho_{solvent})$$

where $\rho_{core}$, $\rho_{corona}$ and $\rho_{solvent}$ are the scattering length densities of the core, corona, and solvent, respectively, where $\rho_{solvent}$ was fixed as 9.44×$10^6$ $Å^{-2}$. The micelle core consists of N polymers, each with volume $V_{core}$, leading to a core radius $r$ which is treated as an independent fitting parameter. The core radius $r$ was fixed using the radius of gyration values from Guinier analysis.

Core form factor and Gaussian form factor are as follows:

$$\Phi(qr) = \frac{sin(qr)-qrcos(qr)}{(qr)^3}$$

$$P_c(q) = 2[exp(-Z) + Z - 1]/Z^2$$

$$Z = (qR_g)^2$$

The Gaussian random coil polymers, of gyration radius $R_g$ (distinctive from that of Guinier analysis), are uniformly distributed around the spherical core, centered at a distance $r + dR_g$ from the micelle center, where $d$ is the factor of non-penetration of Gaussian chains. $V_{corona}$ is defined for each coil.

Core to corona and coil to coil cross terms are approximated as follows:

$$S_{sc}(q) = \Phi(qr)\psi(Z)\frac{sin(q(r+dR_g))}{q(r+dR_g)}$$

$$S_{cc}(q) = \psi(Z)^2\left[\frac{sin(q(r+dR_g))}{q(r+dR_g)}\right]^2$$

$$\psi(Z) = \frac{1-e^{-Z}}{Z}$$

Supplementary Table 1. Amino acid sequence of mdIDPs and GFP variants.

<table>
<tr><th></th><th>net charge at pH 7.4</th><th>length (residue)</th><th>$M_w$</th></tr>
<tr><td rowspan="2">ELPS$_0$-H5$_{+47M}$</td><td>+47.2</td><td>116</td><td>12,578.9 (12,447.7*)</td></tr>
<tr><td colspan="3">MGHHHHHHGGASGVGASGSFRLAKSDKAKRSPGKKKKAVRRSTSPKKAARPRKARSPAKKPKA<br>TARKARKKSRASPKKAKKPKTVKAKSRKASKAKKVKRSKPRAKSGARKSPKKK</td></tr>
<tr><td rowspan="2">ELPS$_{12}$-H5$_{+47M}$</td><td>+47.2</td><td>184</td><td>18,169.9 (18,038.7*)</td></tr>
<tr><td colspan="3">MGHHHHHHSSGGVPGSGVPGSGVPGSGVPGSGVPGSGVPGSGVPGSGVPGSGVPGSGVPGSGVP<br>GSGVPGSGYSWPGGASGVGASGSFRLAKSDKAKRSPGKKKKAVRRSTSPKKAARPRKARSPAKK<br>PKATARKARKKSRASPKKAKKPKTVKAKSRKASKAKKVKRSKPRAKSGARKSPKKK</td></tr>
<tr><td rowspan="2">ELPS$_{24}$-H5$_{+47M}$</td><td>+47.2</td><td>244</td><td>22,939.1 (22,807.9*)</td></tr>
<tr><td colspan="3">MGHHHHHHSSGGVPGSGVPGSGVPGSGVPGSGVPGSGVPGSGVPGSGVPGSGVPGSGVPGSGVP<br>GSGVPGSGVPGSGVPGSGVPGSGVPGSGVPGSGVPGSGVPGSGVPGSGVPGSGVPGSGVPGSGVP<br>GSGYSWPGGASGVGASGSFRLAKSDKAKRSPGKKKKAVRRSTSPKKAARPRKARSPAKKPKATA<br>RKARKKSRASPKKAKKPKTVKAKSRKASKAKKVKRSKPRAKSGARKSPKKK</td></tr>
<tr><td rowspan="2">ELPS$_{48}$-H5$_{+47M}$</td><td>+47.2</td><td>364</td><td>32,477.4 (32,346.2*)</td></tr>
<tr><td colspan="3">MGHHHHHHSSGGVPGSGVPGSGVPGSGVPGSGVPGSGVPGSGVPGSGVPGSGVPGSGVPGSGVP<br>GSGVPGSGVPGSGVPGSGVPGSGVPGSGVPGSGVPGSGVPGSGVPGSGVPGSGVPGSGVPGSGVP<br>GSGVPGSGVPGSGVPGSGVPGSGVPGSGVPGSGVPGSGVPGSGVPGSGVPGSGVPGSGVPGSGVP<br>GSGVPGSGVPGSGVPGSGVPGSGVPGSGVPGSGVPGSGVPGSGVPGSGVPGSGVPGSGYSWPGG<br>ASGVGASGSFRLAKSDKAKRSPGKKKKAVRRSTSPKKAARPRKARSPAKKPKATARKARKKSRA<br>SPKKAKKPKTVKAKSRKASKAKKVKRSKPRAKSGARKSPKKK</td></tr>
<tr><td rowspan="2">ELPS$_{96}$-H5$_{+47M}$</td><td>+47.2</td><td>604</td><td>51,554.0 (51,422.8*)</td></tr>
<tr><td colspan="3">MGHHHHHHSSGGVPGSGVPGSGVPGSGVPGSGVPGSGVPGSGVPGSGVPGSGVPGSGVPGSGVP<br>GSGVPGSGVPGSGVPGSGVPGSGVPGSGVPGSGVPGSGVPGSGVPGSGVPGSGVPGSGVPGSGVP<br>GSGVPGSGVPGSGVPGSGVPGSGVPGSGVPGSGVPGSGVPGSGVPGSGVPGSGVPGSGVPGSGVP<br>GSGVPGSGVPGSGVPGSGVPGSGVPGSGVPGSGVPGSGVPGSGVPGSGVPGSGVPGSGVPGSGVP<br>GSGVPGSGVPGSGVPGSGVPGSGVPGSGVPGSGVPGSGVPGSGVPGSGVPGSGVPGSGVPGSGVP<br>GSGVPGSGVPGSGVPGSGVPGSGVPGSGVPGSGVPGSGVPGSGVPGSGVPGSGVPGSGVPGSGVP<br>GSGVPGSGVPGSGVPGSGVPGSGVPGSGVPGSGVPGSGVPGSGVPGSGVPGSGVPGSGVPGSGVP<br>GSGVPGSGVPGSGVPGSGVPGSGVPGSGVPGSGVPGSGYSWPGGASGVGASGSFRLAKSDKAKR<br>SPGKKKKAVRRSTSPKKAARPRKARSPAKKPKATARKARKKSRASPKKAKKPKTVKAKSRKASK<br>AKKVKRSKPRAKSGARKSPKKK</td></tr>
<tr><td rowspan="2">ELPS$_0$-H5$_{+30L}$</td><td>+30.2</td><td>116</td><td>11,946.3 (11,815.1*)</td></tr>
<tr><td colspan="3">MGHHHHHHGGASGVGASGSFRLAQSDKAGRSPGKSKQAVRRSTSPKGAARPRKARSPASKPQAT<br>ARKARGKSRASPSKAQKPGTVKASSRKASQAKGVKRSSPRAKSGARQSPKGK</td></tr>
<tr><td rowspan="2">ELPS$_{12}$-H5$_{+30L}$</td><td>+30.2</td><td>184</td><td>17,537.3 (17,406.1*)</td></tr>
<tr><td colspan="3">MGHHHHHHSSGGVPGSGVPGSGVPGSGVPGSGVPGSGVPGSGVPGSGVPGSGVPGSGVPGSGVP<br>GSGVPGSGYSWPGGASGVGASGSFRLAQSDKAGRSPGKSKQAVRRSTSPKGAARPRKARSPASKP<br>QATARKARGKSRASPSKAQKPGTVKASSRKASQAKGVKRSSPRAKSGARQSPKGK</td></tr>
<tr><td rowspan="2">ELPS$_{24}$-H5$_{+30L}$</td><td>+30.2</td><td>244</td><td>22,306.5 (22,175.3*)</td></tr>
<tr><td colspan="3">MGHHHHHHSSGGVPGSGVPGSGVPGSGVPGSGVPGSGVPGSGVPGSGVPGSGVPGSGVPGSGVP<br>GSGVPGSGVPGSGVPGSGVPGSGVPGSGVPGSGVPGSGVPGSGVPGSGVPGSGVPGSGVPGSGVP</td></tr>
</table>

| | net charge at pH 7.4 | length (residue) | $M_w$ |
|---|---|---|---|
| | GSGYSWPGGASGVGASGSFRLAQSDKAGRSPGKSKQAVRRSTSPKGAARPRKARSPASKPQATARKARGKSRASPSKAQKPGTVKASSRKASQAKGVKRSSPRAKSGARQSPKGK | | |
| ELPS$_{48}$-H5$_{+30L}$ | +30.2 | 364 | 31,844.8 (31,713.6*) |
| | MGHHHHHHSSGGVPGSGVPGSGVPGSGVPGSGVPGSGVPGSGVPGSGVPGSGVPGSGVPGSGVPGSGVPGSGVPGSGVPGSGVPGSGVPGSGVPGSGVPGSGVPGSGVPGSGVPGSGVPGSGVPGSGVPGSGVPGSGVPGSGVPGSGVPGSGVPGSGVPGSGVPGSGVPGSGVPGSGVPGSGVPGSGVPGSGVPGSGVPGSGVPGSGVPGSGVPGSGVPGSGVPGSGVPGSGVPGSGVPGSGVPGSGVPGSGVPGSGVPGSGYSWPGGASGVGASGSFRLAQSDKAGRSPGKSKQAVRRSTSPKGAARPRKARSPASKPQATARKARGKSRASPSKAQKPGTVKASSRKASQAKGVKRSSPRAKSGARQSPKGK | | |
| ELPS$_{0}$-H5$_{+30H}$ | +30.2 | 73 | 7,938.4 (7,807.2*) |
| | MGHHHHHHGSASGSASKPKATARKARKKSRASPKKAKKPKTVKAKSRKASKAKKVKRSKPRAKSGARKSPKKK | | |
| ELPS$_{12}$-H5$_{+30H}$ | +30.2 | 137 | 13,227.2 (13,096.0*) |
| | MGHHHHHHSSGGVPGSGVPGSGVPGSGVPGSGVPGSGVPGSGVPGSGVPGSGVPGSGVPGSGVPGSGVPGSGYSWPGSASKPKATARKARKKSRASPKKAKKPKTVKAKSRKASKAKKVKRSKPRAKSGARKSPKKK | | |
| ELPS$_{24}$-H5$_{+30H}$ | +30.2 | 197 | 17,996.3 (17,865.1*) |
| | MGHHHHHHSSGGVPGSGVPGSGVPGSGVPGSGVPGSGVPGSGVPGSGVPGSGVPGSGVPGSGVPGSGVPGSGVPGSGVPGSGVPGSGVPGSGVPGSGVPGSGVPGSGVPGSGVPGSGVPGSGVPGSGVPGSGYSWPGSASKPKATARKARKKSRASPKKAKKPKTVKAKSRKASKAKKVKRSKPRAKSGARKSPKKK | | |
| ELPS$_{48}$-H5$_{+30H}$ | +30.2 | 317 | 27,534.7 (27,403.5*) |
| | MGHHHHHHSSGGVPGSGVPGSGVPGSGVPGSGVPGSGVPGSGVPGSGVPGSGVPGSGVPGSGVPGSGVPGSGVPGSGVPGSGVPGSGVPGSGVPGSGVPGSGVPGSGVPGSGVPGSGVPGSGVPGSGVPGSGVPGSGVPGSGVPGSGVPGSGVPGSGVPGSGVPGSGVPGSGVPGSGVPGSGVPGSGVPGSGVPGSGVPGSGVPGSGVPGSGVPGSGVPGSGVPGSGVPGSGVPGSGVPGSGVPGSGVPGSGVPGSGVPGSGYSWPGSASKPKATARKARKKSRASPKKAKKPKTVKAKSRKASKAKKVKRSKPRAKSGARKSPKKK | | |
| ELPS$_{48}$-H5$_{+60M}$ | +60.2 | 395 | 35,797.3 (35,666.1*) |
| | MGHHHHHHSSGGVPGSGVPGSGVPGSGVPGSGVPGSGVPGSGVPGSGVPGSGVPGSGVPGSGVPGSGVPGSGVPGSGVPGSGVPGSGVPGSGVPGSGVPGSGVPGSGVPGSGVPGSGVPGSGVPGSGVPGSGVPGSGVPGSGVPGSGVPGSGVPGSGVPGSGVPGSGVPGSGVPGSGVPGSGVPGSGVPGSGVPGSGVPGSGVPGSGVPGSGVPGSGVPGSGVPGSGVPGSGVPGSGVPGSGVPGSGVPGSGVPGSGVPGSGYSWPGSASASKRSPGKKKKAVRRSTSPKKAARPRKGSASGVGASGSFRLAKSDKAKRSPGKKKKAVRRSTSPKKAARPRKARSPAKKPKATARKARKKSRASPKKAKKPKTVKAKSRKASKAKKVKRSKPRAKSGARKSPKKK | | |
| ELPS$_{48}$-H5$_{+60H}$ | +60.2 | 380 | 34,299.9 (34,168.7*) |
| | MGHHHHHHSSGGVPGSGVPGSGVPGSGVPGSGVPGSGVPGSGVPGSGVPGSGVPGSGVPGSGVPGSGVPGSGVPGSGVPGSGVPGSGVPGSGVPGSGVPGSGVPGSGVPGSGVPGSGVPGSGVPGSGVPGSGVPGSGVPGSGVPGSGVPGSGVPGSGVPGSGVPGSGVPGSGVPGSGVPGSGVPGSGVPGSGVPGSGVPGSGVPGSGVPGSGVPGSGVPGSGVPGSGVPGSGVPGSGVPGSGVPGSGVPGSGVPGSGVPGSGYSWPGSASASKPKATARKARKKSRASPKKAKKPKTVKAKSRKASKAKKVKRSKPRAKSGARKSPKKKGSASKPKATARKARKKSRASPKKAKKPKTVKAKSRKASKAKKVKRSKPRAKSGARKSPKKK | | |
| ELPS$_{48}$-H5$_{+94M}$ | +94.2 | 474 | 44,245.4 (44,114.2*) |

| | net charge at pH 7.4 | length (residue) | $M_w$ |
|---|---|---|---|
| | MGHHHHHHSSGGVPGSGVPGSGVPGSGVPGSGVPGSGVPGSGVPGSGVPGSGVPGSGVPGSGVP<br>GSGVPGSGVPGSGVPGSGVPGSGVPGSGVPGSGVPGSGVPGSGVPGSGVPGSGVPGSGVPGSGVP<br>GSGVPGSGVPGSGVPGSGVPGSGVPGSGVPGSGVPGSGVPGSGVPGSGVPGSGVPGSGVPGSGVP<br>GSGVPGSGVPGSGVPGSGVPGSGVPGSGVPGSGVPGSGVPGSGVPGSGVPGSGVPGSGYSWPGSA<br>SASGVGASGSFRLAKSDKAKRSPGKKKKAVRRSTSPKKAARPRKARSPAKKPKATARKARKKSR<br>ASPKKAKKPKTVKAKSRKASKAKKVKRSKPRAKSGARKSPKKKGSASGVGASGSFRLAKSDKA<br>KRSPGKKKKAVRRSTSPKKAARPRKARSPAKKPKATARKARKKSRASPKKAKKPKTVKAKSRKA<br>SKAKKVKRSKPRAKSGARKSPKKK | | |
| GFP-12 | -11.4 | 248 | 27,775.4 (27,644.2*) |
| | MGHHHHHHGGASKGEELFTGVVPILVELDGDVNGHKFSVRGEGEGDATEGKLTLKFICTTGKLPV<br>PWPTLVTTLTYGVQCFSRYPDHMKQHDFFKSAMPEGYVQERTISFKDDGTYKTRAEVKFEGDTLV<br>NRIELKGIDFKEDGNILGHKLEYNFNSHNVYITADKQENGIKANFKIRHNVEDGSVQLADHYQQN<br>TPIGDGPVLLPDNHYLSTQSALSKDPNEDRDHMVLLEFVTAAGITHGMDELYK | | |
| GFP-18 | -17.4 | 248 | 27,765.2 (27,634.0*) |
| | MGHHHHHHGGASKGEELFTGVVPILVELDGDVNGHKFSVRGEGEGDATEGKLTLKFICTTGELPV<br>PWPTLVTTLTYGVQCFSRYPDHMDQHDFFKSAMPEGYVQERTISFKDDGTYKTRAEVKFEGDTLV<br>NRIELKGIDFKEDGNILGHKLEYNFNSHDVYITADKQENGIKANFKIRHNVEDGSVQLADHYQQN<br>TPIGDGPVLLPDNHYLSTESALSKDPNEDRDHMVLLEFVTAAGITHGMDELYK | | |
| GFP-24 | -23.4 | 248 | 27,754.0 (27,622.8*) |
| | MGHHHHHHGGASKGEELFTGVVPILVELDGDVNGHEFSVRGEGEGDATEGKLTLKFICTTGELPV<br>PWPTLVTTLTYGVQCFSDYPDHMDQHDFFKSAMPEGYVQERTISFKDDGTYKTRAEVKFEGDTL<br>VNRIELKGIDFKEDGNILGHKLEYNFNSHDVYITADKQENGIKAEFKIRHNVEDGSVQLADHYQQ<br>NTPIGDGPVLLPDNHYLSTESALSKDPNEDRDHMVLLEFVTAAGIDHGMDELYK | | |
| Dark GFP-24 | -23.4 | 248 | 27,621.8 (27,490.6*) |
| | MGHHHHHHGGASKGEELFTGVVPILVELDGDVNGHEFSVRGEGEGDATEGKLTLKFICTTGELPV<br>PWPTLVTTLGGGVQCFSDYPDHMDQHDFFKSAMPEGYVQERTISFKDDGTYKTRAEVKFEGDTL<br>VNRIELKGIDFKEDGNILGHKLEYNFNSHDVYITADKQENGIKAEFKIRHNVEDGSVQLADHYQQ<br>NTPIGDGPVLLPDNHYLSTESALSKDPNEDRDHMVLLEFVTAAGIDHGMDELYK | | |
| GFP-30 | -29.4 | 248 | 27,770.9 (27,639.7*) |
| | MGHHHHHHGGASKGEELFDGVVPILVELDGDVNGHEFSVRGEGEGDATEGELTLKFICTTGELPV<br>PWPTLVTTLTYGVQCFSDYPDHMDQHDFFKSAMPEGYVQERTISFKDDGTYKTRAEVKFEGDTL<br>VNRIELKGIDFKEDGNILGHKLEYNFNSHDVYITADKQENGIKAEFEIRHNVEDGSVQLADHYQQ<br>NTPIGDGPVLLPDDHYLSTESALSKDPNEDRDHMVLLEFVTAAGIDHGMDELYK | | |
| mTurquoise2 | -13.5 | 250 | 27,950.7 (27,818.5*) |
| | MGHHHHHHGGAVSKGEELFTGVVPILVELDGDVNGHKFSVSGEGEGDATYGKLTLKFICTTGELP<br>VPWPTLVTTLSWGVQCFARYPDHMKQHDFFKSAMPEGYVQERTIFFKDDGNYKTRAEVKFEGDT<br>LVNRIELKGIDFKEDGNILGHKLEYNYFSDNVYITADKQENGIKANFKIRHNIEDGGVQLADHYQQ<br>NTPIGDGPVLLPDNHYLSTQSKLSKDPNEDRDHMVLLEFVTAAGITLGMDELYK | | |

**Supplementary Table 1. Amino acid sequence of mdIDPs and FP variants.** Neutral domains and cationic domains on the sequences are labelled as gray and blue, respectively. FP sequences are labeled in green. Peptide linkers and affinity purification tags are in black. *Expected molecular weights were determined assuming cleavage of the initial methionine residue.

### Supplementary Figures

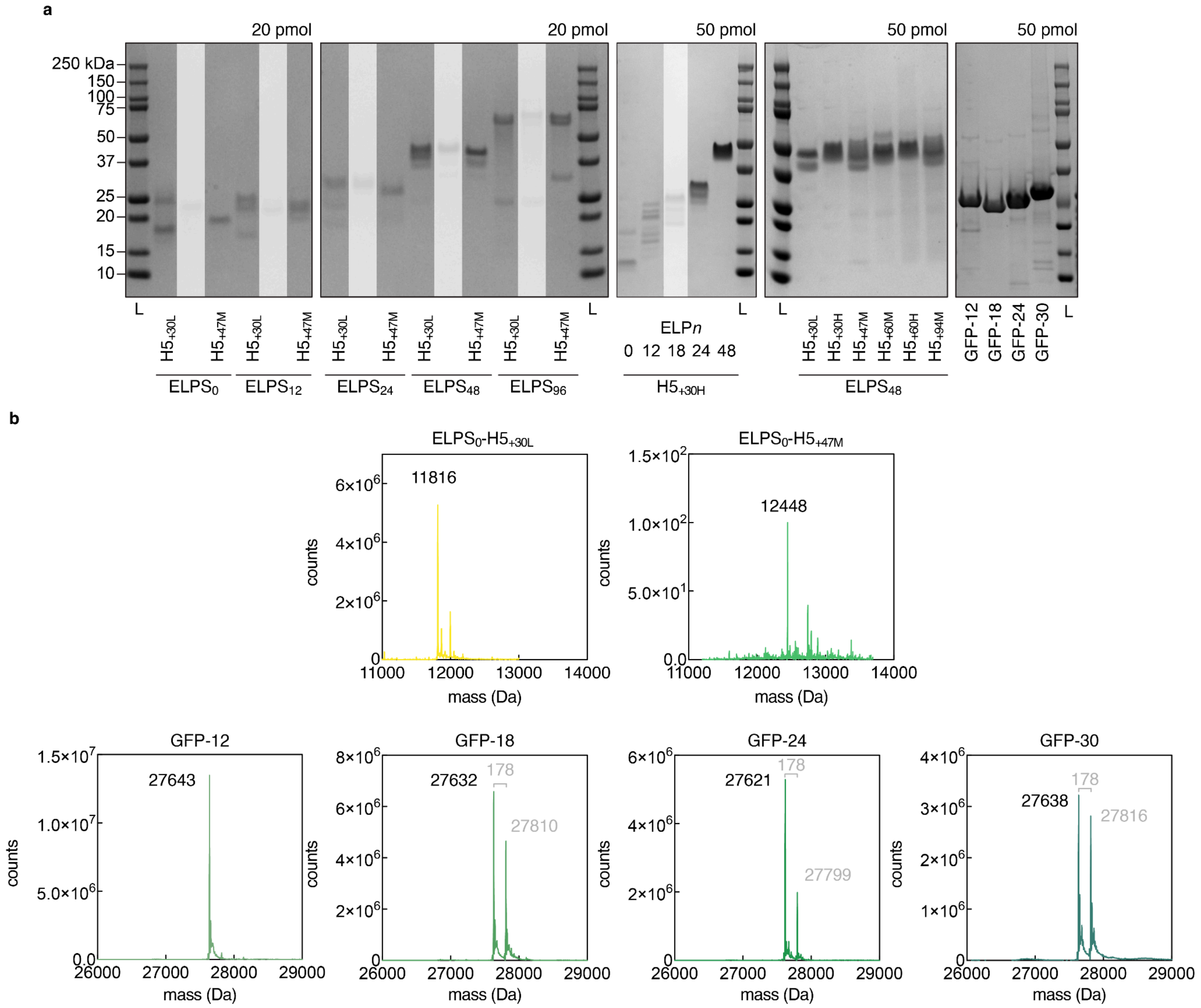

**Supplementary Figure 1. Analysis of purified proteins.** (a) SDS-PAGE analysis of mdIDPs and GFP variants. 20 pmol or 50 pmol of the proteins were loaded per well for SDS-PAGE analysis. Shaded lanes correspond to proteins or fractions that were not used in subsequent experiments. (b) Mass spectrometry analysis of H5 and GFP variants. All expressed proteins that could be analyzed by LC-MS have the correct expected molecular weight. Expected molecular weights were determined assuming cleavage of the initial methionine residue. The 178 Da shift, labelled in gray, from the expected molecular weight observed in the GFP samples, corresponds to a-N-6-Phosphogluconoylation of a His-tag in *E. coli.*[12]

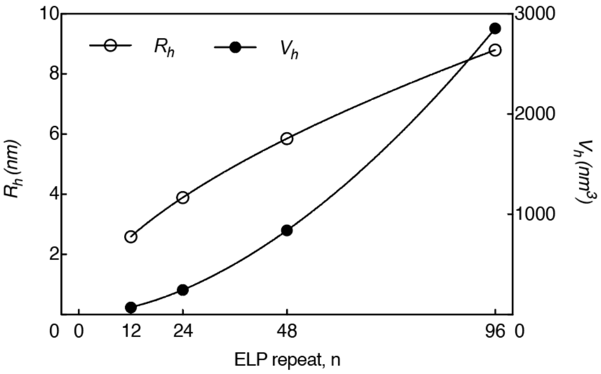

**Supplementary Figure 2. Hydrodynamic radius and hydrodynamic volume of ELP variants.** $R_h$ and $V_h$ values were estimated using the equations and assumptions below.[13,14] The radius of gyration $R_g$ can be estimated by the Flory Scaling Law (1)

$$R_g = aN^{\nu} \quad (1)$$

where $a$ is the monomer length (Kuhn length, 0.35 nm for average amino acid residue), $N$ is the number of monomers, and ν is the Flory exponent (0.588 for good solvent and flexible chain). For disordered and flexible chains, the ratio of the radius of gyration $R_g$ to the hydrodynamic radius $R_h$ is approximately 1.5. The hydrodynamic volume $V_h$ can be calculated as:

$$V_h = \frac{4}{3}\pi R_h^{3} \quad (2)$$

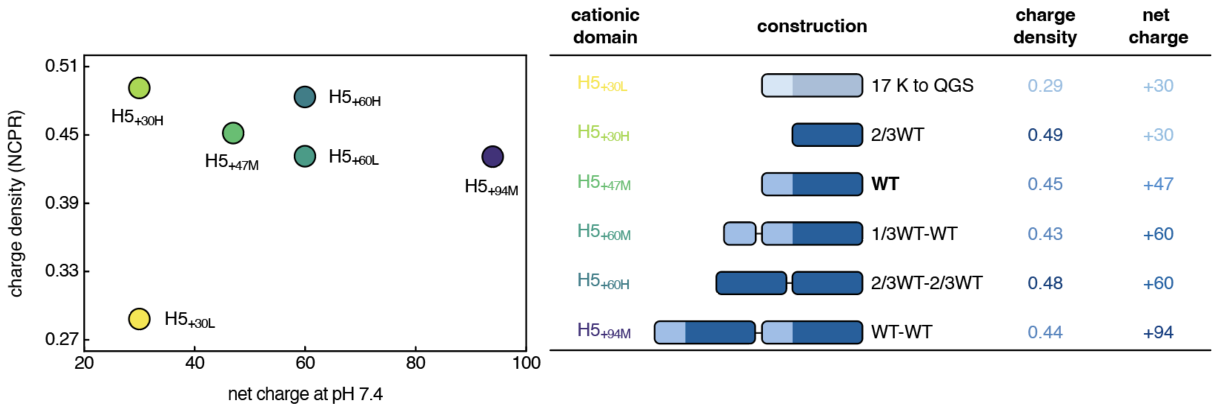

**Supplementary Figure 3. Net Charge, charge density, and construction of cationic domains.** The cationic domain was derived from the disordered region of the nucleosome linker histone H5 (PDB ID: 7XVM), specifically residues 87–190, which were selected based on their high cationic charge density and predicted disorder. The wild-type sequence of this region ($H5_{+47M}$) carries a net charge of +47 at pH 7.4, with a charge density of 0.45 (calculated as the ratio of cationic residues to total residues), and is predicted to be highly disordered with an IUPred3A score of 0.99. To systematically vary the electrostatic properties, several variants were designed. The $H5_{+30L}$ variant was generated by substituting 17 lysines in the $H5_{+47M}$ sequence with uncharged residues (glutamine, glycine, or serine), resulting in a net charge of +30 and a reduced charge density of 0.29. The $H5_{+30H}$ was designed by selecting the C-terminal 57 residues of the $H5_{+47M}$ sequence, which have a higher charge density of 0.49. The $H5_{+60M}$ variant was created by appending a charge-sparse region to the $H5_{+47M}$ sequence, yielding a net charge of +60 and a charge density of 0.43. The $H5_{+60H}$ variant was constructed by repeating the $H5_{+30H}$ segment, resulting in a charge density of 0.48, while the $H5_{+94M}$ variant was formed by duplicating the $H5_{+47M}$ sequence, with a final charge density of 0.44. The charge densities of the repeated variants are slightly lower than the shorter unit sequences due to the presence of neutral linkers introduced between repeated segments.

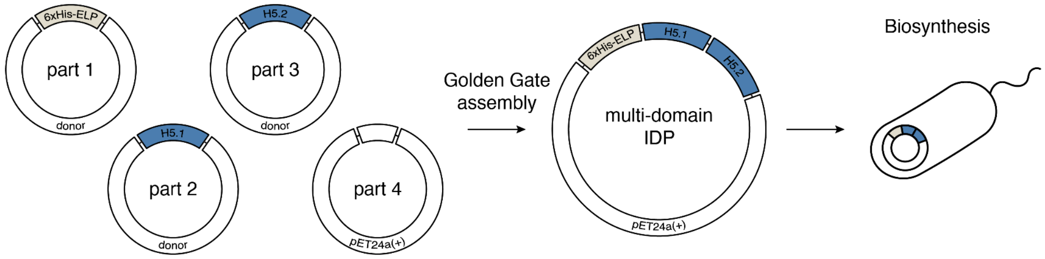

| Part 1 | Part 2 | Part 3 | Part 4 | Final Assembly |
|---|---|---|---|---|
| 6xHis-ELPS$_0$ | H5 (17 K to QGS) | | pET24a(+) backbone | ELPS$_n$-H5$_{+30L}$ |
| 6xHis-ELPS$_{12}$ | H5 (2/3WT) | | | ELPS$_n$-H5$_{+30H}$ |
| 6xHis-ELPS$_{18}$ | H5 (WT) | | | ELPS$_n$-H5$_{+47M}$ |
| 6xHis-ELPS$_{24}$ | H5 (1/3WT) | H5 (WT) | | ELPS$_n$-H5$_{+60M}$ |
| 6xHis-ELPS$_{48}$ | H5 (2/3WT) | H5 (2/3WT) | | ELPS$_n$-H5$_{+60H}$ |
| 6xHis-ELPS$_{96}$ | H5 (WT) | H5 (WT) | | ELPS$_n$-H5$_{+94M}$ |

**Supplementary Figure 4. Four-part Golden Gate library for biosynthesis of mdIDPs.** Variations in neutral domain length, and cationic domain charge were modularly encoded and assembled as part of a Golden Gate library. Each module was flanked by defined junction sequences to enable directional ligation to adjacent parts. Different parts were stored in donor vectors containing the coding regions of gene fragments of interest, along with BsaI recognition sites and defined junctions. The pET24a(+) vector backbone provides the start and stop codons flanking the upstream end of part 1 and the downstream end of part 4, respectively. Each part is joined via BsaI restriction sites that introduce defined junctions between modules during assembly. Example mdIDP constructs prepared using the Golden Gate library include the assembly of H5$_{+13L}$ with H5$_{+47M}$, yielding the H5$_{+60M}$ variant; two tandem repeats of H5$_{+30H}$ yielding H5$_{+60H}$; and two tandem repeats of H5$_{+47M}$ yielding the H5$_{+94M}$ variant. A total of 72 mdIDP constructs were generated through combinatorial assembly from the constructed library.

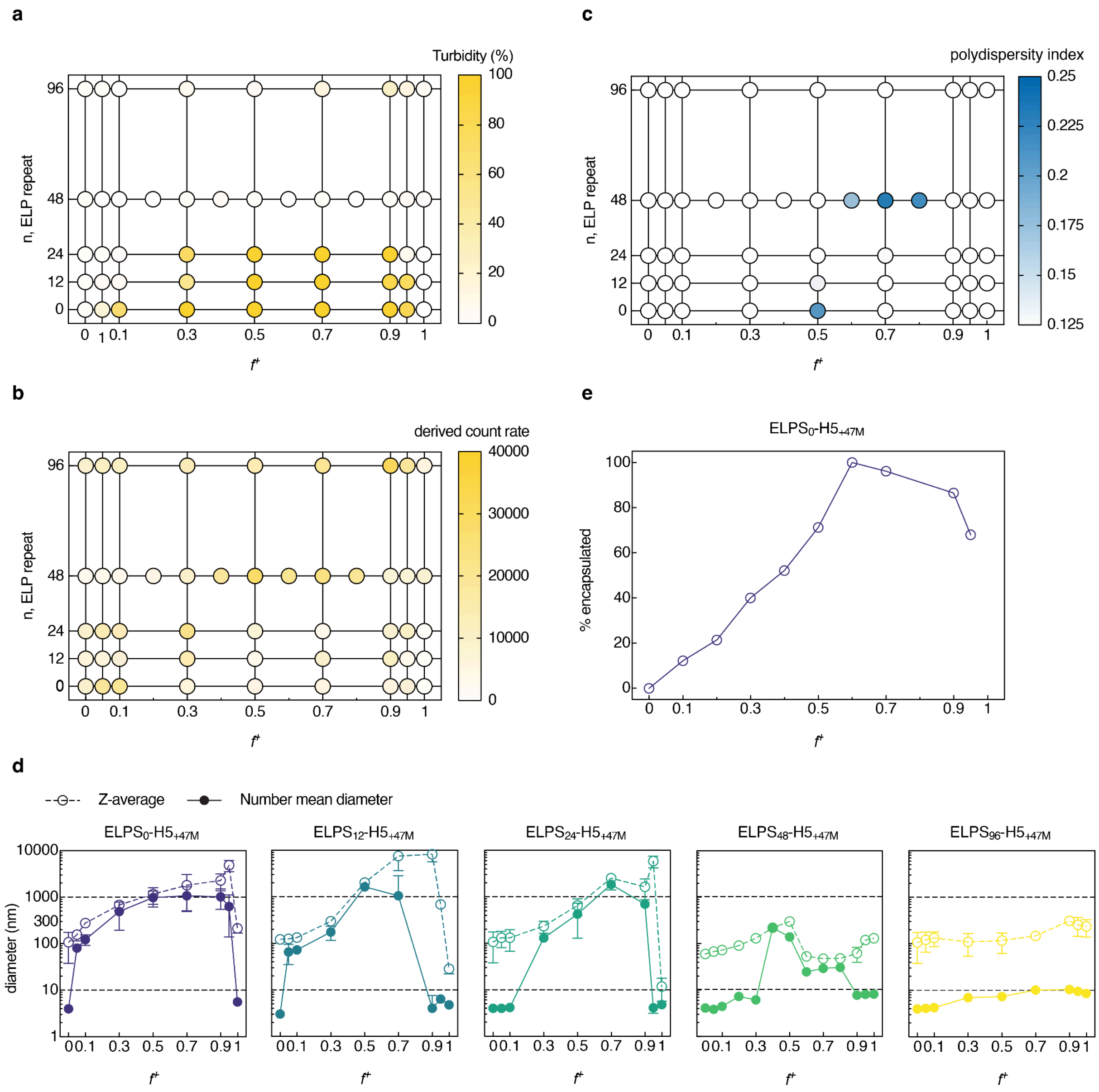

**Supplementary Figure 5. Characterization of assemblies with ELPs of varying length.** ELPS$_n$-H5$_{+47M}$ and GFP-24 were mixed at varied charge fractions. The assemblies were characterized by (a) turbidity analysis and (b-d) DLS. The DLS (b) derived count rate, (c) polydispersity index (PDI), and (d) Z-average and number mean diameter are shown as a function of the mixing ratio ($f^+$) and ELP length. (e) GFP encapsulation in the coacervate phase with the ELP$_0$-H5$_{+47M}$ variant. The total macromolecule concentration was fixed at 100 μM for experiments performed in (a-d) and 50 μM for the encapsulation assay in (e).

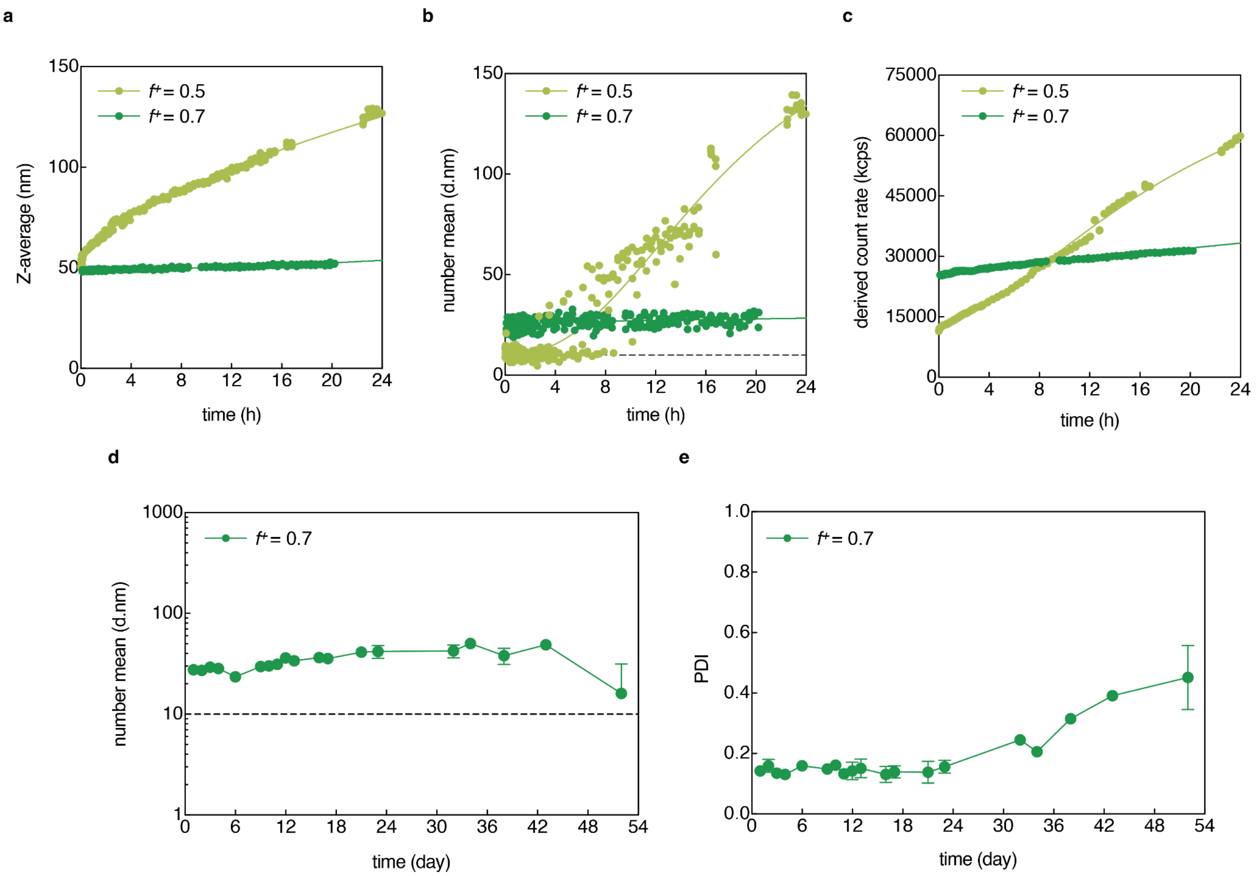

**Supplementary Figure 6. Metastable nanoassembly and micellar nanoparticle assembly kinetics.** (a) Z-average, (b) number mean diameter, and (c) derived count rate by DLS. $ELPS_{48}$-$H5_{+47M}$ and GFP-24 were mixed at $f^+ = 0.5$ and $f^+ = 0.7$ to represent metastable nanoassembly and micellar nanoparticle assembly, respectively. The total macromolecule concentration was fixed at 100 µM. (d) Number mean diameter and (e) polydispersity index of micellar nanoparticles over time from DLS analysis. Assemblies were prepared using $ELPS_{48}$-$H5_{+47M}$ and GFP-24. Micellar nanoparticles in (d–e) were stored at 4 °C, and DLS analysis was performed at 25 °C.

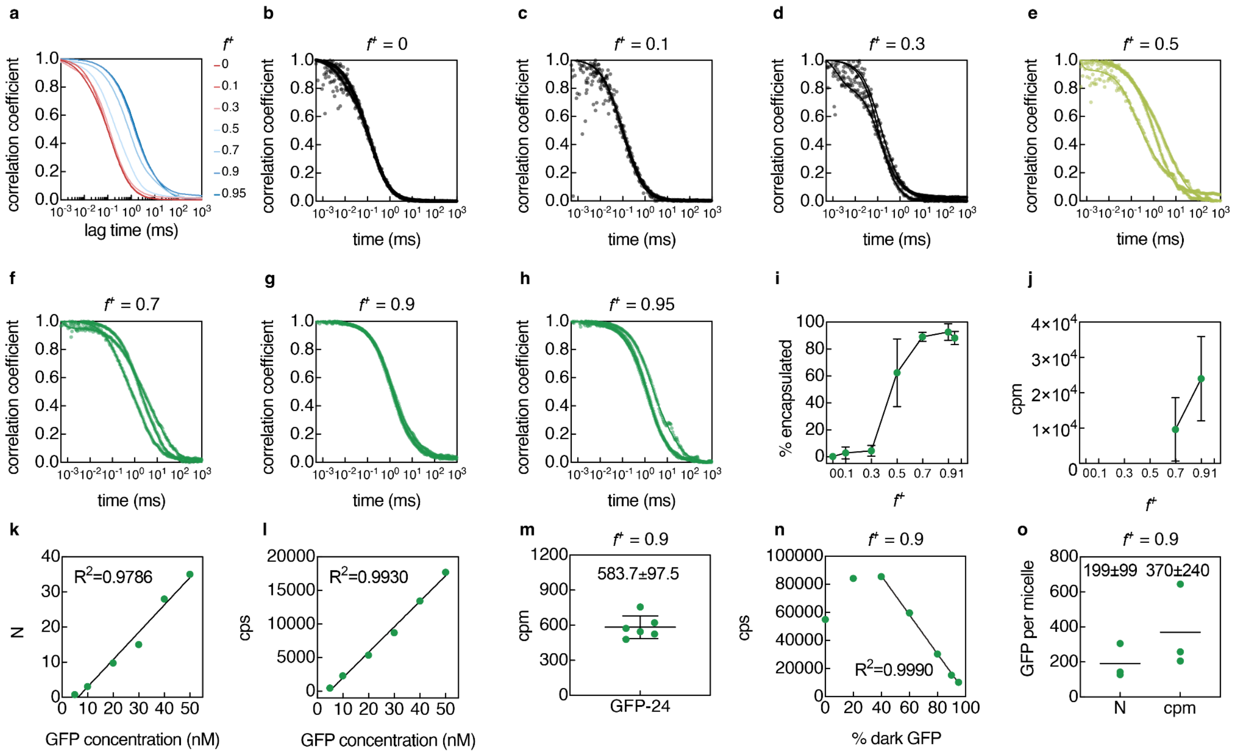

**Supplementary Figure 7. Characterization of assemblies by FCS.** The assemblies were prepared from mixtures of ELPS$_{48}$-H5$_{+47M}$ and GFP-24. (a) Mean correlation curves from three replicates as a function of mixing ratio. Individual correlation curves from replicates (n=3) of (b) $f^+ = 0$, (c) $f^+ = 0.1$, (d) $f^+ = 0.3$, (e) $f^+ = 0.5$, (f) $f^+ = 0.7$, (g) $f^+ = 0.9$, and (h) $f^+ = 0.95$. (i) GFP encapsulation in nanoscale assemblies with ELPS$_{48}$-H5$_{+47M}$, equivalent to Figure 2c. (j) Counts per molecule (cpm) of $f^+ = 0.7$ and $f^+ = 0.9$ assemblies. (k) Number of molecules ($N$) and (l) counts per second (cps) of GFP-24 at different concentrations. Both cps and N showed a linear correlation at the given conditions. (m) Counts per molecule (cpm) of GFP-24 determined by measurements from (k) and (l). (n) cps of particles at $f^+ = 0.9$ mixing ratio of ELPS$_{48}$-H5$_{+47M}$ and GFP-24 as a function of the % of dark GFP(-24). (o) Estimates of GFP-24 per particle at $f^+ = 0.9$, determined using either the measured N or cpm. Triplicate measurements used to estimate the number of GFP-24 per particle using the number of molecules (N) were collected on three separate days. Triplicate measurements used to estimate the number of GFP-24 per particle using the cpm were collected on the same day using the same measured cpm of free GFP-24 and calibration setting.

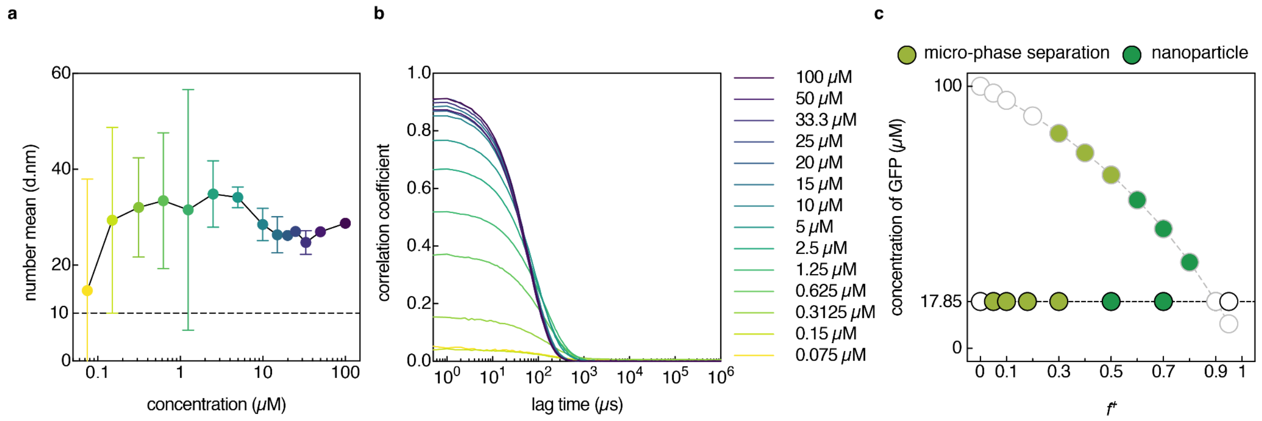

**Supplementary Figure 8. Micellar nanoparticle formation at different concentrations.** (a) Number mean diameter analysis and (b) correlation curves of nanoparticles following serial dilution. Assemblies were prepared with ELPS$_{48}$-H5$_{+47M}$ (54.4 µM) and GFP-24 (45.6 µM) for a total macromolecule concentration of 100 µM and $f^{+}$= 0.7. The micellar nanoparticles were then serially diluted and measured by DLS in (a–b) were assembled at $f^{+}$= 0.7. (c) Phase diagram showing metastable nanoassembly and micellar nanoparticle assembly at a fixed GFP concentration of 17.85 µM in comparison to constant macromolecule concentration and changing GFP concentration, outlined in gray.

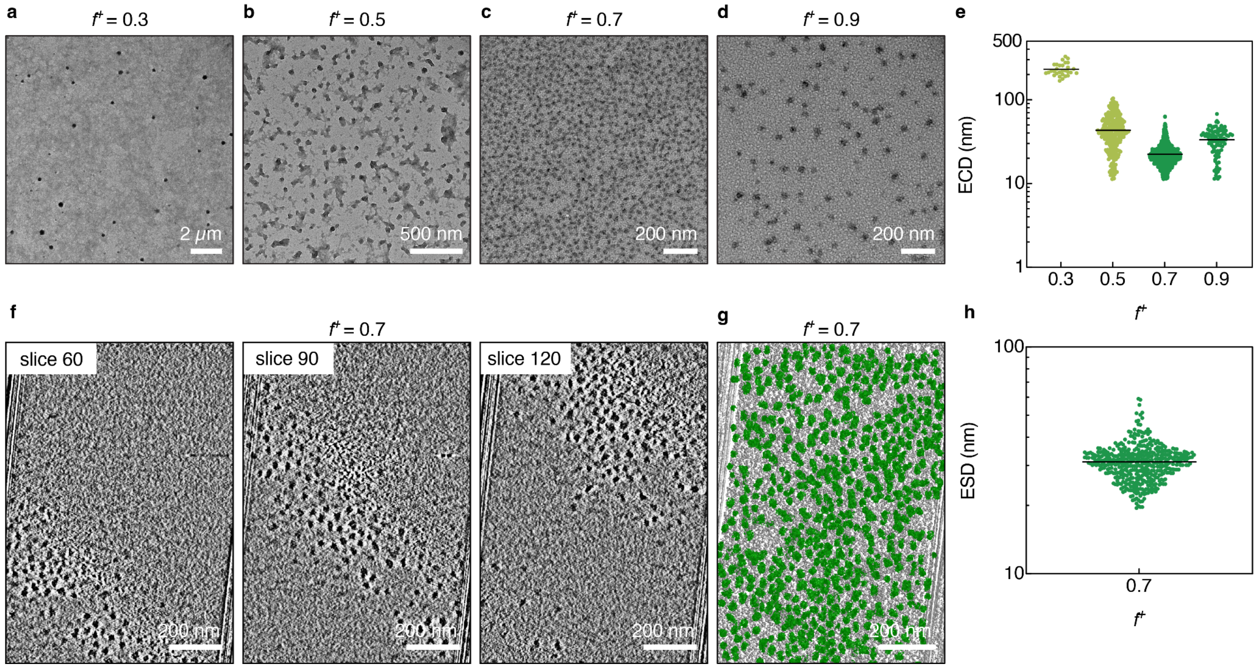

**Supplementary Figure 9. Size characterization of metastable nanoassemblies and nanoparticles by TEM and cryo-ET.** The assemblies were generated by mixing $ELPS_{48}$-$H5_{+47M}$ and GFP-24. TEM images of the assemblies at (a) $f^+ = 0.3$, (b) $f^+ = 0.5$, (c) $f^+ = 0.7$, and (d) $f^+ = 0.9$. (e) Assembly diameter from images (a), (b), (c), and (d) were quantified by approximating the particle area and converting it to an equivalent circular diameter (ECD). (f) Cryo-ET images of $f^+ = 0.7$ nanoparticles are shown across multiple Z slices of the reconstructed tomogram. (g) The full tomogram corresponding to (f) if shown segmented to highlight individual nanoparticles. (h) Micellar nanoparticle diameter from image (g) was quantified by approximating particle volume and converting it to an equivalent spherical diameter (ESD). See Supplementary Movie 1 for full tomogram.

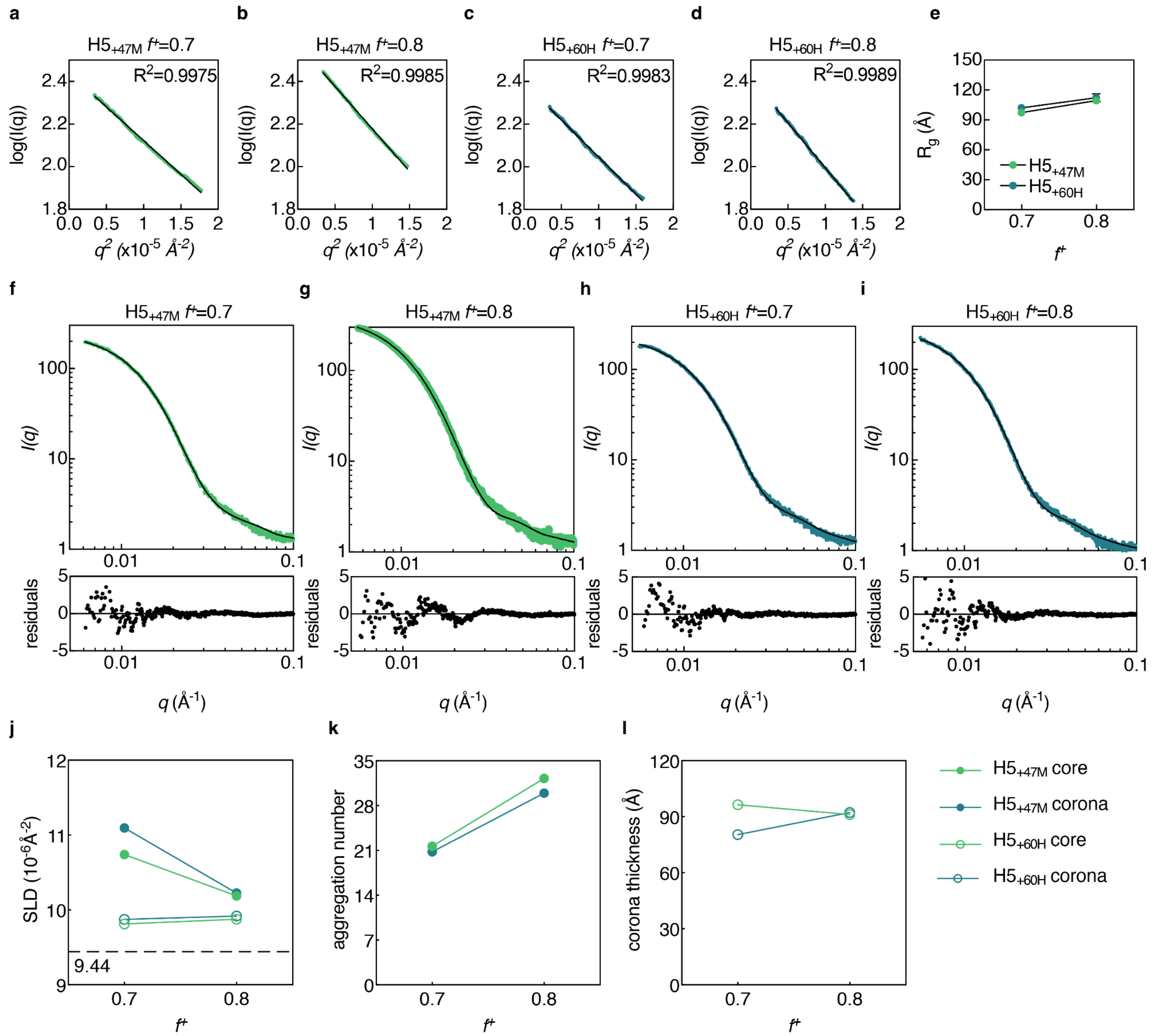

**Supplementary Figure 10. SAXS analysis of protein nanoparticles.** 1D scattering profile in the Guinier region and corresponding Guinier fits for nanoparticles formed with GFP-24 and (a) ELPS$_{48}$-H5$_{+48M}$ $f^+$ = 0.7, (b) $f^+$ = 0.8, (c) ELPS$_{48}$-H5$_{+60H}$ $f^+$ = 0.7, and (d) $f^+$ = 0.8. (e) Radius of gyration ($R_g$) of nanoparticles obtained from Guinier analysis. Polymer-micelle model form factor fits to the 1D scattering profiles and corresponding residuals for (f) ELPS$_{48}$-H5$_{+47M}$ $f^+$ = 0.7, (g) $f^+$ = 0.8, (h) ELPS$_{48}$-H5$_{+60H}$ $f^+$ = 0.7, and (i) $f^+$ = 0.8 nanoparticles with GFP-24. (j) Scattering length density (SLD), (k) aggregation number, and (l) corona thickness, obtained from form factor fitting of the micelles formed at two different charge fractions with the two different mdIDPs and GFP-24.

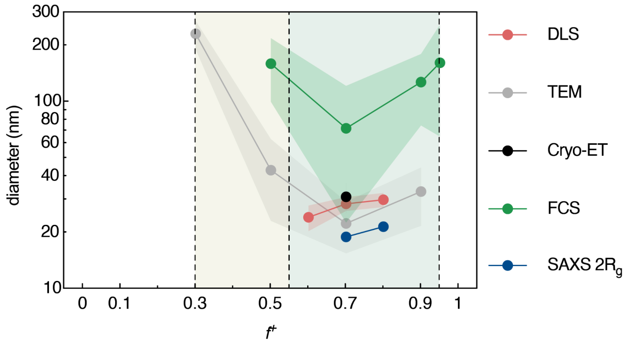

**Supplementary Figure 11. Size characterization of assemblies using various techniques.** The assemblies were formed from $ELPS_{48}$-$H5_{+47M}$ and GFP-24 at various charge fractions. Micellar nanoparticles form at charge fractions ranging from $f^+ = 0.6$ to 0.9 (denoted by the green background), whereas metastable nanoassemblies form at $f^+ = 0.3$ to 0.5 (denoted by the olive background). Data points represent the average value and shaded areas denote the standard deviation.

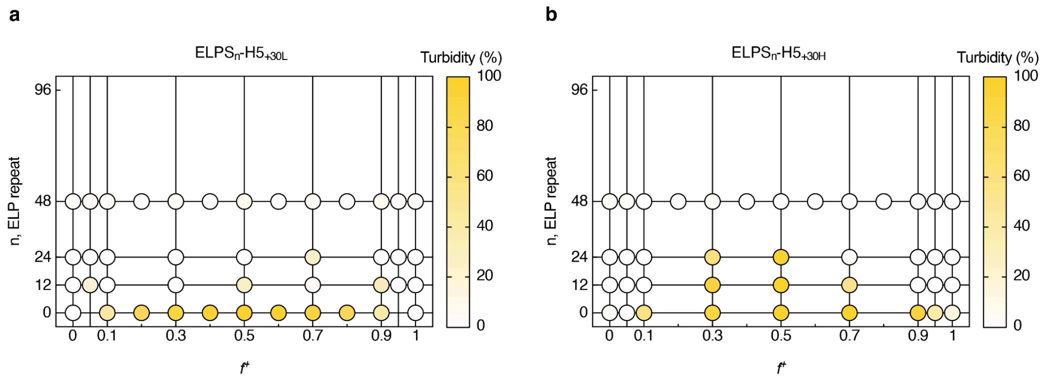

**Supplementary Figure 12. Turbidity of (a) ELPS$_n$-H5$_{+30L}$, and (b) ELPS$_n$-H5$_{+30H}$ with different lengths of neutral domains (n) and GFP-24.** Turbidity persisted for H5$_{+30H}$ under charge-neutral conditions to ELPS$_{24}$, whereas H5$_{+30L}$ exhibited no notable turbidity for any ELP length above zero.

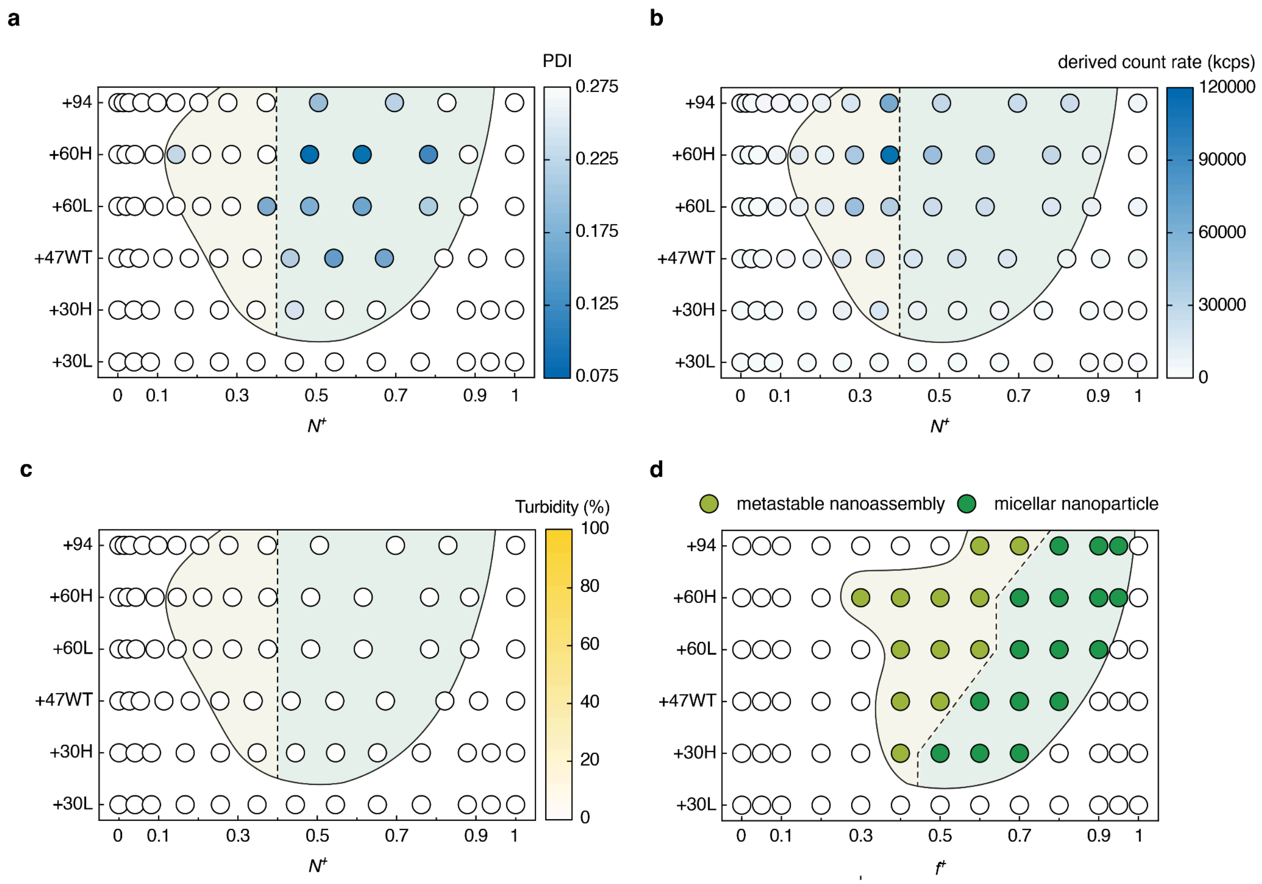

**Supplementary Figure 13. Phase diagram illustrating metastable nanoassembly and micellar nanoparticle assembly regime for engineered mdIDPs with varying cationic domains.** The neutral domain and anionic globular protein were fixed as $ELPS_{48}$ and GFP-24, respectively. Heat map of (a) polydispersity index (PDI) and (b) derived count rate obtained from DLS measurements as a function of the stoichiometric mixing ratio ($N^+$). (c) Turbidity measurements indicated assemblies were too small to scatter visible light. The vertical dotted lines in (a), (b) and (c) indicate $N^+ = 0.4$, where $N^+$ is the molar fraction of the cationic species. (d) Phase diagram illustrating metastable nanoassembly and micellar nanoparticle assembly regime plotted with charge fraction, $f^+$, as a variable for mixing ratio.

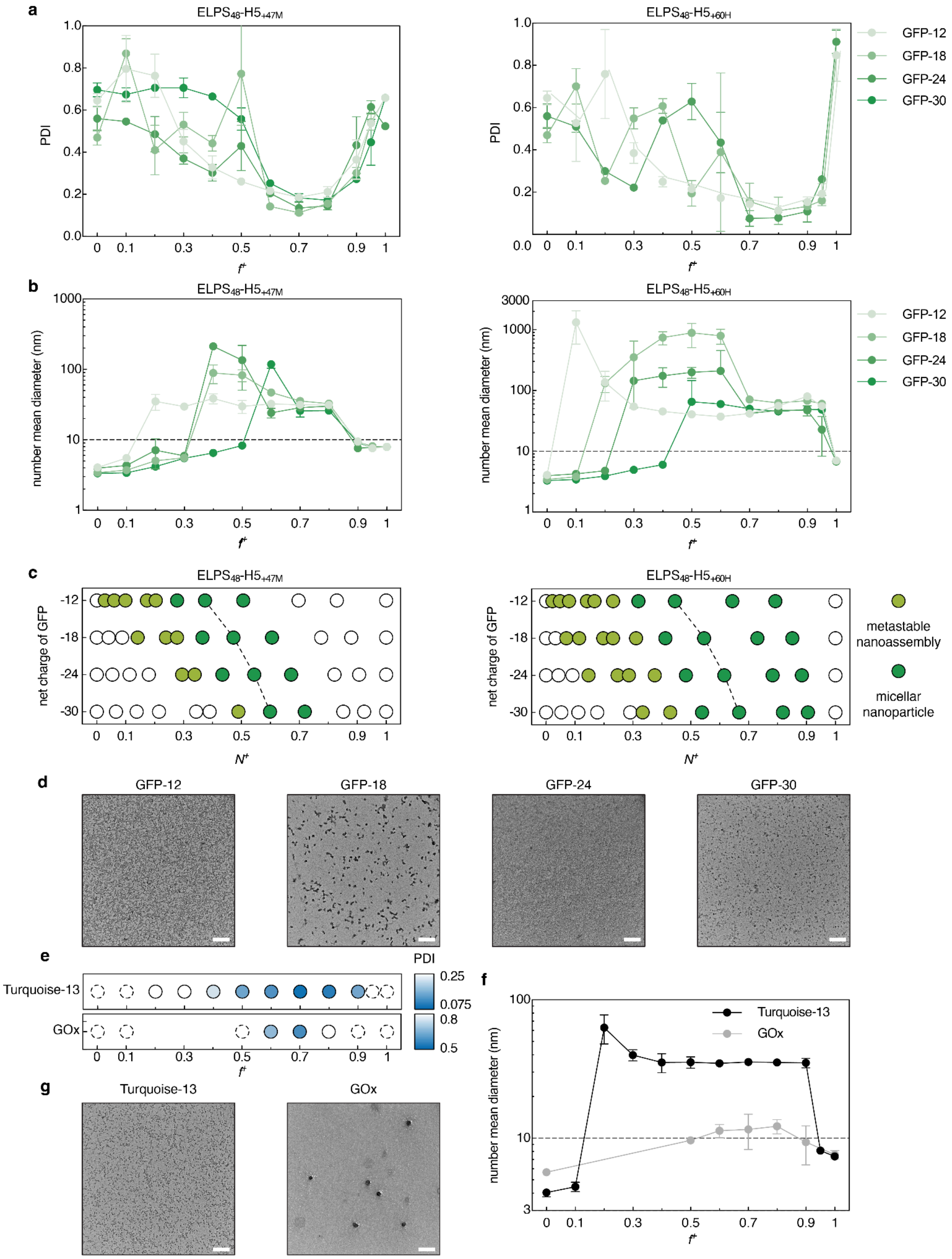

**Supplementary Figure 14. Micellar nanoparticle formation with various globular protein cargos.** (a) PDI and (b) number mean diameter of assemblies from (left) $ELPS_{48}$-$H5_{+47M}$ and (right) $ELPS_{48}$-$H5_{+60H}$ with GFPs of various net charge. (c) Phase diagram displaying metastable nanoassembly and micellar nanoparticle assembly as a function of stoichiometric mixing ratio for (left) $ELPS_{48}$-$H5_{+47M}$ and (right) $ELPS_{48}$-$H5_{+60H}$. (d) TEM image of micellar nanoparticles assembled from $ELPS_{48}$-$H5_{+47M}$ with GFPs of various net charge at $f^{+}= 0.7$. (e) PDI and (f) number mean diameter of assemblies from $ELPS_{48}$-$H5_{+47M}$ with mTurquoise2 and GOx. For (e), only PDI values corresponding to the metastable nanoassembly and micellar nanoparticle assembly conditions are shown; regions outside the nanoparticle assembly regime are indicated with dotted circles. TEM images of nanoparticles assembled from $ELPS_{48}$-$H5_{+47M}$ with (g) (left) mTurquoise2 and (right) GOx at $f^{+}$= 0.7. Scale bars: 500 nm.
